# Enhanced Omicron subvariant cross-neutralization efficacy of a monovalent SARS-CoV-2 BA.4/5 mRNA vaccine encoding a noncleaved, nonfusogenic spike antigen

**DOI:** 10.1101/2023.09.10.557088

**Authors:** Han Young Seo, Haewon Jung, Hawon Woo, Hae-Gwang Jung, Hee Cho, Yeonju Bak, Se-Young Lee, Yu-Min Son, Gone Yoon, Seo-Yeon Hwang, Inho Park, Jeon-Soo Shin, Jong-Won Oh

## Abstract

The rapid emergence of diverse SARS-CoV-2 variants, notably the Omicron variant, poses challenges to vaccine development. Here, we present a noncleaved, nonfusogenic spike (S) protein eliciting robust B- and T-cell immune responses against Omicron BA.5. The antigen incorporates the R685S and R815A mutations, effectively preventing the shedding of the S1 subunit and eliminating fusogenic activity of the resulting S antigen, termed S(SA). Through reverse genetic analysis, we found that the noncleaved form S protein with the R685S mutation enhances ACE2-dependent viral entry in vitro compared to the wild-type S protein, without increasing the virulence of the mutant virus in mice. The mRNA vaccine encoding the Omicron BA.4/5 S(SA) antigen conferred protective immunity in mice following two doses of 1 μg Ψ-UTP- or UTP-incorporated mRNA vaccines. Despite a roughly 6-fold reduction in neutralizing potency, both mRNA vaccines exhibited broad neutralizing efficacy against Omicron subvariants, including the XBB lineage variants XBB.1.5 and XBB.1.16.

## Introduction

The ongoing global threat of COVID-19, caused by SARS-CoV-2, remains substantial. The extended duration of this pandemic can be attributed to the virus’s propensity to mutate, leading to the emergence of immune evasive variants. Of particular concern are mutations within the immunodominant spike (S) protein, which can result in the evasion of pre-existing immunity. One notable variant, the SARS-CoV-2 Omicron variant, has garnered significant attention since its emergence in late 2021. Characterized by an unprecedented number of mutations within the *S* gene, this variant has the capacity to almost completely escape the pre-existing immune system^1–5^.

The S protein plays a pivotal role in SARS-CoV-2 entry into host cells, mediating receptor binding and membrane fusion^6^. Proteolytic cleavage of the S protein by cellular proteases at the S1/S2 and S2′ cleavage sites is essential for viral entry into host cells. The S1/S2 processing site is located between the receptor-binding S1 subunit and the membrane fusion-mediating S2 subunit. The other cleavage site (S2′) is located immediately upstream of the fusion peptide within the S2 subunit. Cleavage at both sites occurs sequentially and is of great importance during priming in SARS-CoV and MERS CoV infection^7,8^.

SARS-CoV-2 employs the angiotensin-converting enzyme 2 (ACE2)-dependent transmembrane protease serine subtype 2 (TMPRSS2)-mediated cellular membrane fusion and alternative endosomal protease-mediated fusion pathways for S protein activation and cellular entry^6^. Notably, the characteristic polybasic furin cleavage site (FCS) at the S1/S2 site, which is not found in any other lineage B beta-coronaviruses^9^, confers improved pre-priming efficiency, thereby contributing to its enhanced infectivity^10^. Pre-priming at the S1/S2 site during S protein biosynthesis results in S1 and S2 subunit cleavage, followed by priming at the S2ʹ site upon receptor binding, leading to the exposure of the fusion peptide^6^. Hoffmann et al. demonstrated the critical role of the SARS-CoV-2 FCS in infecting human lung cells, a finding enhancing our understanding of the pathogenicity of SARS-CoV-2^11^. However, this FCS is highly prone to deletions and mutations^12–14^, raising questions about its clinical implications and potential effects on vaccine antigen design.

The mRNA-1273 and BNT162b2 are two approved mRNA vaccines for COVID-19^15,16^. These vaccines encode the complete SARS-CoV-2 S protein sequence with two proline mutations (K986P and V987P, referred to as “2P”) that allow the antigen to be locked in a prefusion state^17^. Although the 2P-mutated S antigen is proven to be an effective antigen for SARS-CoV-2 vaccines, the antigen is still prone to cleavage by furin proprotein convertase during its biogenesis^16,18^. The S1 fragment released after the pre-priming processing can activate proinflammatory innate immune responses through the TLR4 signaling pathway^19^. The clinical implications of proteolytic cleavage and shedding of the detached vaccine antigen remain to be fully defined, although Ogata et al. observed the shedding of a significant amount of the S1 subunit after mRNA-1273 vaccination^20^.

In this study, we devised a nonfusogenic, noncleaved form of the S antigen through functional analysis of S variants that emerged during the passage of SARS-CoV-2 in TMPRSS2-deficient VeroE6 cells. We demonstrate that the FCS-defective S protein with an R685S substitution [S(R685S)], which conferred viral growth advantage in ACE2-expressing cells, exhibited reduced fusogenecity. By introducing an additional mutation at the S2ʹ cleavage site (R815A), we generated a fusion incompetent, noncleaved form of the S protein and evaluated its immunogenicity using Ψ-UTP- or UTP-incorporated mRNA vaccines. Our findings demonstrate that this novel S antigen design has the potential to enhance the safety profile of the mRNA vaccine and bolster its efficacy against emerging SARS-CoV-2 variants, offering potential solutions to combat the ongoing global health threat posed by COVID-19.

## Results

### Harnessing the R685S cell-culture adaptive mutation in the S protein for vaccine antigen design

The Omicron BA.4/5 mRNA vaccine developed in this study was founded on the noncleaved form of the S protein. This design stemmed from functional analyses conducted on cell-culture-adapted S variants of S-clade SARS-CoV-2 strains, NCCP43326 and NCCP43331, initially isolated in early 2020 at the Korean Center for Disease Control (KCDC). During the propagation of these S clade SARS-CoV-2 strains, by infecting VeroE6 cells to prepare passage 1 (P1) stocks, it was observed that S protein processing was severely impaired in the NCCP43326 strain. This resulted in a negligible amount of the S2 fragment produced during biogenesis, in contrast to the other strain NCCP43331 (Fig. 1a). Notably, the NCCP43326 strain, predominantly expressing the uncleaved S protein, exhibited faster growth than NCCP43331 in VeroE6 cells at 1 and 2 dpi, while their peak titers at 3 dpi were comparable (Fig. 1b).

**Fig. 1.**
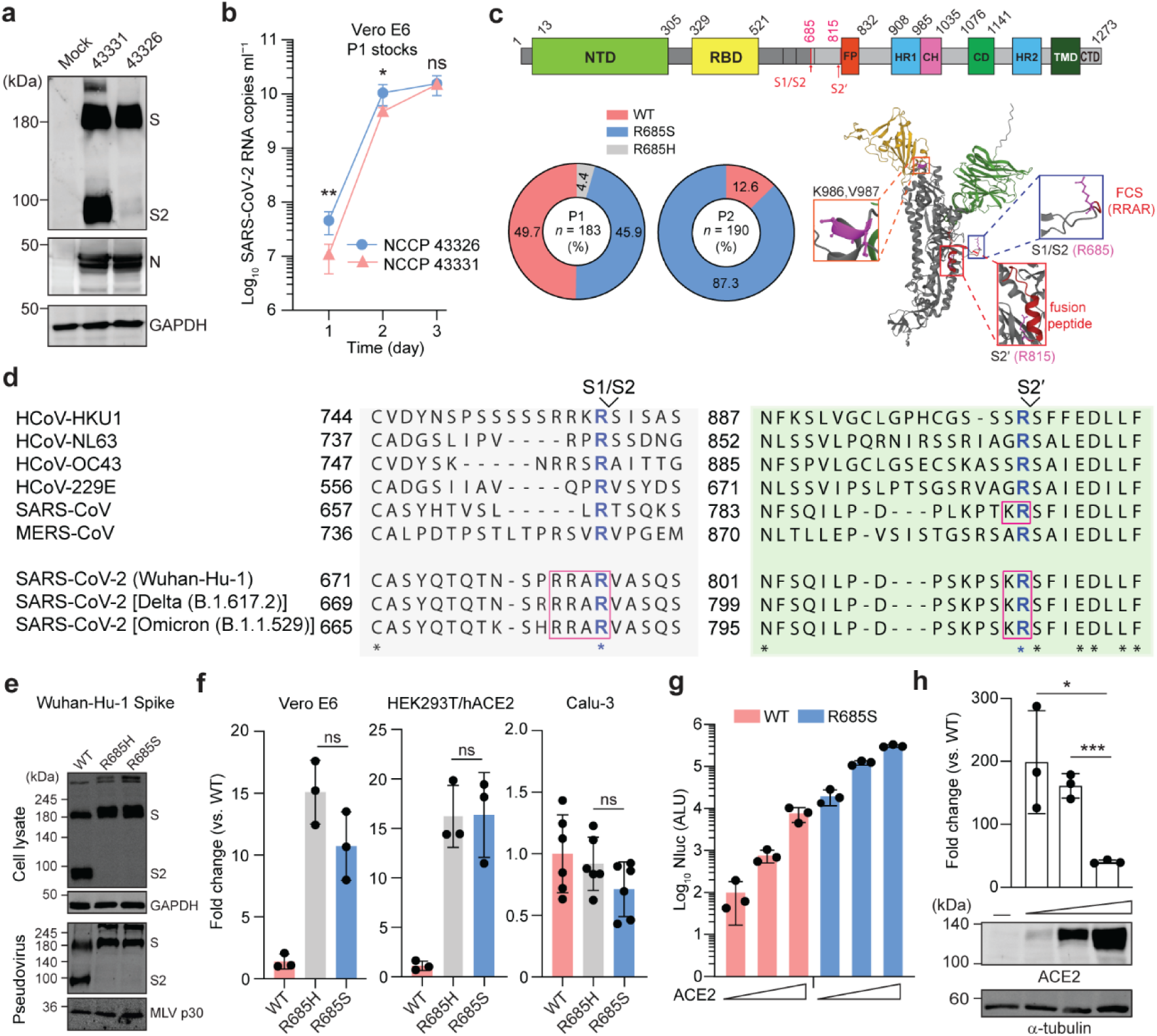
Impact of cell culture-adapted SARS-CoV-2 S protein variants on viral entry via ACE2 receptor. **a**, Differential processing of S protein in VeroE6 cells infected with indicated S-clade SARS-CoV-2 viral stocks (MOI = 0.01), assessed through immunoblotting using an S2 domain-specific monoclonal antibody at 2 dpi. N, nucleocapsid protein. **b**, Viral growth kinetics in VeroE6 cells infected with NCCP43326 or NCCP43331 strain P1 stock (MOI = 0.01). Results show mean ± SD from two independent experiments with biological triplicates in each experiment. **c**, A schematic of SARS-CoV-2 S protein domains with proteolytic cleavage sites, S1/S2 and S2′. Profile of S protein variants with various R685 mutations in the P1 and P2 stocks of NCCP43326 strain. The R685 site at the conserved “RRAR” motif (blue box) is indicated on the S protein structure, along with the R815 residue (red box) at the S2′ site. The K986 and V987 residues mutated to prolines in the approved mRNA vaccines are presented in the orange box. **d**, Amino acid sequence variations at the two processing sites, S1/S2 and S2′, among seven distinct human pathogenic coronaviruses. **e**, Furin-cleavage resistant S variants with an R685H or R685S substitution, loaded onto MLV pseudoviruses and in HEK293T cells. GAPDH and MLV p30, loading controls for cell lysates and pseudoviruses, respectively. **f**, Impact of R685 mutations on entry of SARS-CoV-2 S-pseudotyped MLVs into various cell lines. Results depict mean (fold change) ± SD from three or six biological replicates. **g**,**h**, Influence of hACE2 expression levels on S(R685S)-pseudotyped MLV entry into HEK293T cells. Shown in (**h**) are the entry fold changes between WT S- and S(R685S)-loaded pseudotype MLVs and the immunoblots showing hACE2 expression levels. Statistical significance was determined by unpaired student t-test of the log10-transformed RNA copies ml^-1^ at the same time point (b). For (f and h), unpaired ordinary one-way ANOVA with Tukey’s multiple comparisons test was used. \**P* < 0.05; ** *P* < 0.01; *** *P* < 0.001; ns, not significant.

Subsequent RNA sequencing analysis of the NCCP43326 P1 stock unveiled a major proportion of the S variant with an R685S substitution (45.9 %) and a minor read of R685H substitution (4.37 %) out of a total read of 183 (number of contigs covering all 3 nucleotides for the 685th codon) (Fig. 1c, Extended Fig. 1a, Supplementary Table 1, 2). These single amino acid changes occurred at the S1/S2 site, located within the external random coil structure connecting the S1 and S2 subunits, which is recognized by furin-like proteases^21^. Crucially, the S2ʹ cleavage by TMPRSS2 or cathepsin L is a pivotal step for early viral entry at the cellular membrane or for late or slow entry through the endosomal pathway^6^. Remarkably, no cell-culture-adapted mutations emerged at the S2′ cleavage site located upstream of the fusion peptide (FP) (Fig. 1c, top). Further analysis of the S variants in the P2 stock of the NCCP43326 strain revealed that the R685S variant outcompeted R685H, H655Y, and R682W (all at 0%), as well as the original sequence R685 (12.6%) (Fig. 1c, Supplementary Table 1). The two deletion mutants ΔQTQ and ΔQTQTN remained as minor variants in the P2 stock (Supplementary Table 2). Previously, these deletion mutants were also observed in cell-culture-adapted S variants^12–14^. Along with the R685S mutation, the R685H, H655Y, and the two deletion mutants indeed exhibited varying levels of S1/S2 cleavage efficiency in HEK293T cells (Extended Data Fig. 1b).

The distinctive “RRAR” polybasic motif found at the S1/S2 site is a sequence unique to SARS-CoV-2, and the last arginine residue is highly conserved among human pathogenic coronaviruses (Fig. 1d). This residue is highly conserved across more than 13 million patient-isolated SARS-CoV-2 sequences (Extended Data Fig. 2a), suggesting its crucial role in SARS-CoV-2 S protein biogenesis^21^. Supporting this hypothesis, any modification of this R685 residue led to the inhibition of S1/S2 cleavage (Extended Data Fig. 2b). Additionally, the two basic residues “KR” at the S2′ site are conserved in SARS-CoVs. In particular, the R815 residue exhibited conservation across various human pathogenic CoVs. Notably, variants with mutation at the R815 site constituted only 0.003% of 13,423,789 SARS-CoV-2 sequences as of February 1, 2023 (Extended Data Fig. 2c). Intriguingly, changes in the R815 residue to amino acids other than the R815H and R815K substitutions interfered with S1/S2 cleavage in HEK293T cells (Extended Data Fig. 2d), highlighting that alterations in the S2′ site with non-basic amino acids can hinder the pre-priming of SARS-CoV-2 S protein, potentially affecting its fusogenic activity by inhibiting further processing of the pre-primed S protein. Based on these findings, we aimed to devise a noncleaved form of the S antigen, one that offers enhanced safety by preventing the shedding of the S1 subunit and the initiation of S-mediated cell-to-cell fusion with minimal alteration to the S antigen sequence.

### The S mutant with an R685S substitution enhances ACE2-dependent viral entry while exhibiting reduced fusogenic activity

We investigated the impact of two noncleaved form S variants, carrying an R685S or R685H mutation, on viral entry using these S variant-pseudotyped murine leukemia viruses (MLVs). These variants did not generate the S2 fragment, leading to the generation of pseudoviruses loaded solely with the uncleaved S protein (Fig. 1e). These pseudoviruses more efficiently entered ACE2-expressing VeroE6 and HEK293T/hACE2 cells than wild-type S-pseudotyped MLV (Fig. 1f). However, the entry enhancement was not observed in TMPRSS2-positive human lung epithelial cell line Calu-3. In HEK293T cells with minimal ACE2 expression, ectopic ACE2 expression promoted S(R685S)-pseudotyped virus entry over the S(WT)-pseudotyped virus entry. Entry efficiency increase was particularly notable in cells with lower ACE2 levels (Fig. 1g,h). This indicated the selective advantage of the S(R685S) variant in ACE2-positive, TMPRSS2-negative cells, like VeroE6 cells, which primarily use the endosomal pathway as the major entry route for SARS-CoV-2^22^. This is supported by the insensitivity of VeroE6 cells to the TMRPSS2 inhibitor camostat but sensitivity to the cathepsin L inhibitor E-64d, (Extended Data Fig. 3a,b). A similar drug response was observed in HEK293T/hACE2 cells, but the opposite inhibitory effect was displayed by these drugs in Calu-3, verifying that the TMPRSS2-expressing Calu-3 preferentially utilizes the fast entry pathway via viral membrane fusion at the target cell surface.

In E-64d-sensitive, camostat-resistant HEK293T/hACE2 cells, S(R815A)-pseudotyped virus displayed no detectable virus entry, indicating impaired cathepsin L-mediated S protein processing (Fig. 2a). Additionally, the nullified viral entry observed with the pseudotype virus loaded with the S mutant carrying R685S and R815A substitutions underscored that the R815A mutation decreased the heightened viral entry mediated by R685S to the background level.

**Fig. 2.**
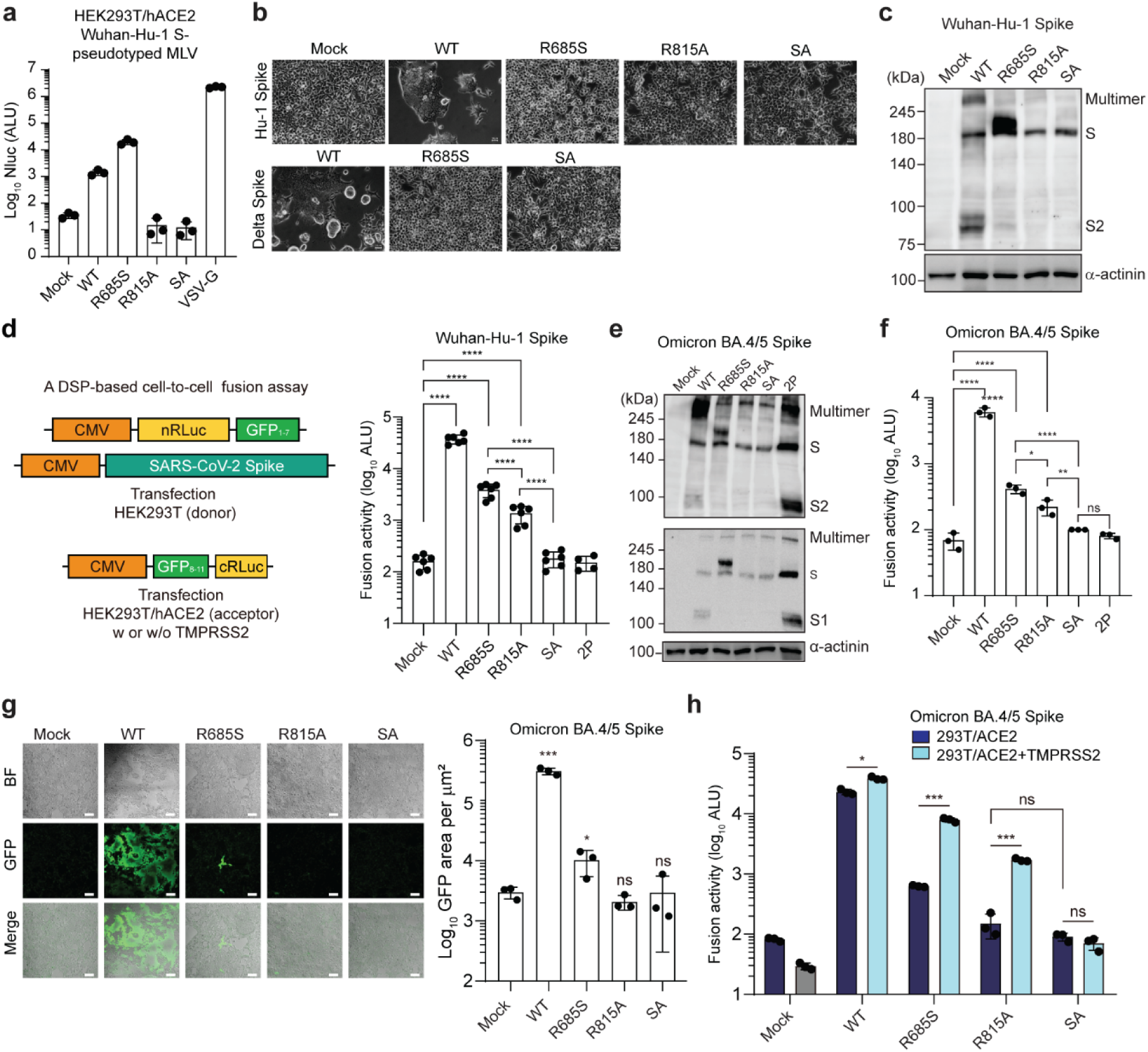
Impact of proteolytic cleavage-resistant S protein variants on viral entry and ACE-dependent S-mediated cell-to-cell fusion. **a**, Pseudotype virus entry assays performed on HEK293T/hACE2 cells. **b**, Cell-to-cell fusion activity evaluated for Wuhan-Hu-1 or Delta S protein and its derivatives in HEK293T/hACE2, assessed 2 days post-transfection by monitoring multinuclear syncytia formation. Scale bar, 50 μm. **c**, Differential processing of Wuhan-Hu-1 S-derived mutant proteins in HEK293T cells, assessed 2 days post-transfection by immunoblotting analysis. **d**, Schematic of the cell-to-cell fusion assay using dual-split proteins (DSP) (left). Fusion activity of Wuhan-Hu-1 S-derived mutant proteins. **e**, Proteolytic cleavage profile of Omicron BA.4/5 S derivatives, analyzed in HEK293T cells by immunoblotting. **f**, Cell-to-cell fusion activity of the BA.4/5 S derivatives with the indicated amino acid changes. **g**, Confocal microscopic analysis of the GFP signal from conjoined DSPs 24 h after co-culture of the acceptor and donor cells. Scale bar, 100 μm. The fusion activity of modified S proteins was accessed by quantification of the GFP-positive area. **h**, Impact of TMPRSS2 expression on ACE2-dependent S protein-mediated cell-cell fusion. In panels (**d,f,g,h**), each data point represents mean ± SD from at least three biological replicates. Statistical significance was determined using one-way ANOVA with Tukey’s multiple comparisons test on log_10_-transformed data. \**P* < 0.05; ** *P* < 0.01; *** *P* < 0.001; **** *P* < 0.001; ns, not significant.

Initial fusogenic activity assessment of these S variants in HEK293T/hACE2 cells revealed that Wuhan-Hu-1 S protein triggered cell-to-cell fusion despite minimal TMPRSS2 expression (Fig. 2b). This suggested the presence of another protease that can trigger the exposure of the fusion-peptide from the S protein through cleavage at the S2ʹ site cleavage following ACE2 binding. Importantly, the cleavage site-mutated S proteins, S(R685S), S(R815A), and S(SA) substantially lost the cell-fusion activity. Wuhan-Hu-1 S(R685S) and S(R815A) mutants generated minimal S2 fragments in HEK293T cells compared to wild-type S (Fig. 2c). Utilizing a DSP-based cell-to-cell fusion assay, we demonstrated Wuhan-Hu-1 S(R685S) and S(R815A) retained residual fusogenic activity. However, S(SA) antigen, like prefusion S(2P), completely lost fusogenic activity (Fig. 2d). S(R685H) also showed reduced fusogenic activity, lower than S(H655Y) and ΔQTQ/ΔQTQTN deletion mutants (Extended Data Fig. 1c), demonstrating that S1/S2 cleavage efficiency correlates with fusion activity. For Omicron BA.4/5 S, S(R685S), S(R815A), and S(SA) mutants displayed similar incompetence in S1/S2 processing and S-ACE2 interaction-mediated cell fusion (Fig. 2e,f). Cell fusion activity of these S variants was not observed when fusion events were quantified based on GFP-signal in the cells fused between S-expressing-donor cells and ACE2-expressing acceptor cells (Fig. 2g).

The fusogenic activity of Omicron BA.4/5 S(R685S) and S(R815A) significantly increased upon TMPRSS2 expression in acceptor HEK293T/hACE2 cells (Fig. 2h). In line with this, only BA.4/5 WT S generated a detectable level of S2ʹ fragments when expressed with hACE2 (Extended Data Fig. 4a). However, when expressed in the HEK293T/hACE2 cells ectopically expressing TMPRSS2, S(R685S) as well as WT S, but not S(R815A) and S(SA) produced substantial amounts of S2′ fragments (Extended Data Fig. 4b). These results highlighted ACE2-binding as a prerequisite for fusion triggered by TMPRSS2-mediated S2′ cleavage. Nevertheless, the S(SA) antigen exhibited no fusogenic activity even with both ACE2 and TMPRSS2, underlining this uncleaved, nonfusogenic S protein as a safe vaccine antigen that avoids shedding the S1 subunit (Fig. 2h).

### Impact of R685S mutation on viral propagation and virulence in mice

To ascertain that the growth and entry enhancement conferred by the R685S mutation can be recapitulated with a homogeneous SARS-CoV-2 strain, we generated two recombinant viruses: one expressing GH-clade WT S protein with the D614G mutation, and the other carrying the R685S mutation. These recombinant viruses were generated using an infectious cDNA clone of the GH clade SARS-CoV-2 YS006 strain^23^ (further details to be reported elsewhere) (Fig. 3a). The virus expressing mutant S, rYS006[S(R685S)], exhibited augmented viral growth compared to the parental strain rYS006 (Fig. 3b), resulting in larger plaques on A549/hACE2 cells (Fig. 3c). In K18-hACE2 mice, survival rates showed no significant difference between recombinant rYS006 and rYS006[S(R685S)] (Fig. 3d,e). Viral RNA copy numbers and infectious titers were comparable between the WT and R685S groups (Fig. 3f,g). Collectively, these findings indicate that S(R685S) did not exhibit heightened pathogenic characteristics in mice, although it facilitated accelerated SARS-CoV-2 propagation in ACE2-positive and TMPRSS2-deficient cells *in vitro*.

**Fig. 3.**
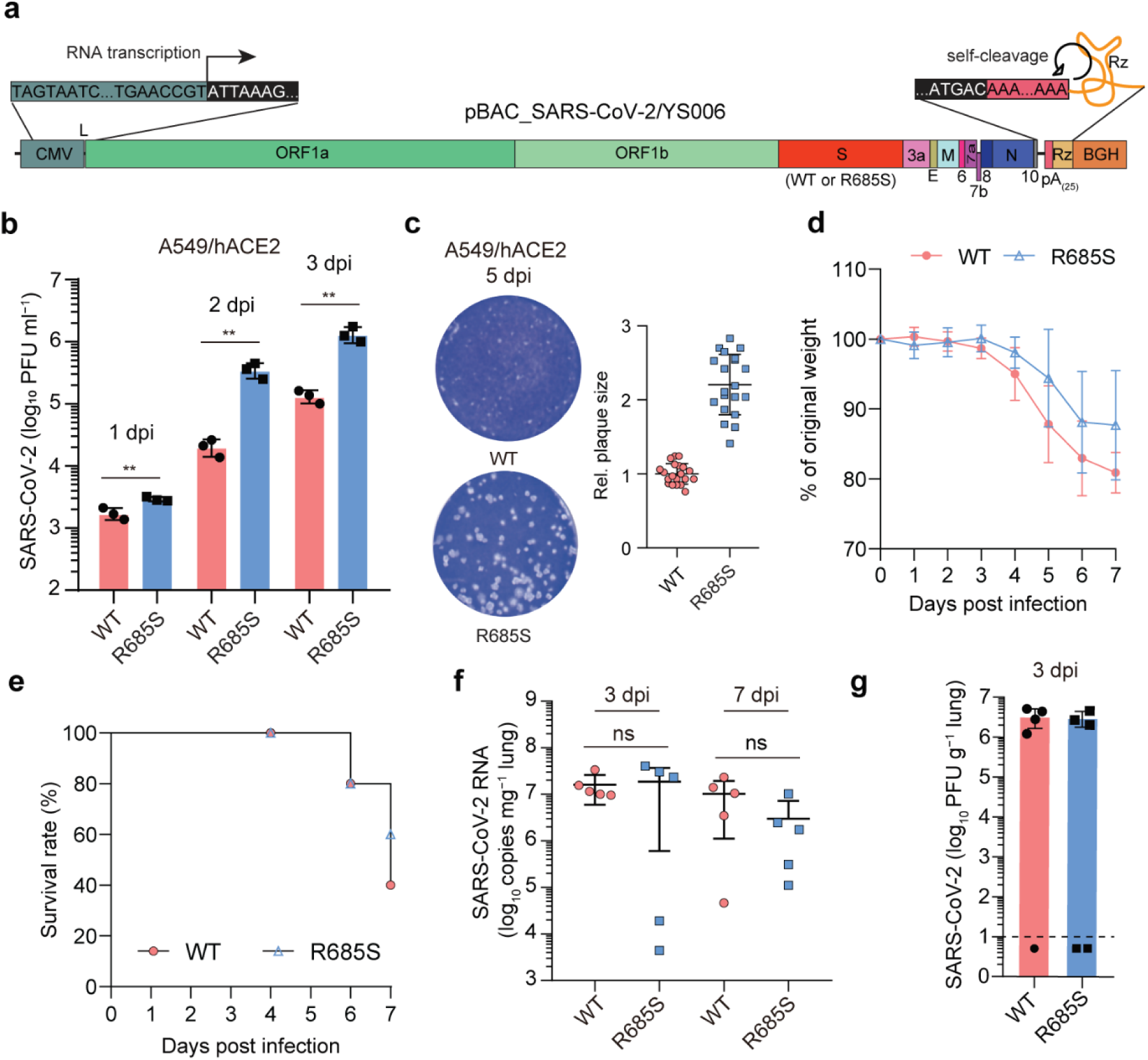
Influence of the R685S mutation on viral growth and pathogenesis in K18/hACE2 mice. **a**, An infectious cDNA clone of a recombinant SARS-CoV-2/YS006 expressing GH-clade S or S(R685S). **b**, Growth kinetics of SARS-CoV-2 rYS006(WT) and rYS006(R685S) variant. Viral loads were assessed following infection of A549/hACE2 with the indicated recombinant viruses at an MOI of 0.01. The results represent the mean ± SD of three biological replicates. **c**, Representative plaque images at 5 dpi of A549/hACE2 cells, showing relative diameters of plaques (*n* = 20 plaques from 3 independent experiments). Plaque diameters were measured digitally using images with a sizing bar. **d**–**g**, Seven-to-eight-week-old K18/hACE2 transgenic mice were intranasally infected with the respective viruses (*n* = 10 per group). **d**, Weight change was monitored for 7 days after infection. **e**, Survival rate was evaluated up to 7 days post-infection. Tissue viral loads in lung homogenates were measured by RT-qPCR (**f**) and plaque assay (**g**). Each data point represents the viral load from two technical replicates for RT-qPCR or two biological replicates for the plaque assay. The dotted line in (**g**) represents the assay limit of detection (LOD). Statistical significance was determined by an unpaired Student *t*-test of the raw data (**d**) or log_10_-transformed data (**b**,**f**). \*\**P* <0.01; ns, not significant.

### Antibody and memory T cell responses induced by BA.5 S(SA)-coding mRNA vaccines produced with Ψ-UTP or UTP

Despite its limited amino acid modifications, the S(SA) antigen, characterized by enhanced ACE2-dependent entry and the lack of cell-to-cell fusion activity, must elicit substantial antibody and T-cell immune responses to be considered a suitable vaccine antigen. We assessed the immunogenicity of the BA.5 S(SA) antigen using the mRNA comprised of 5′UTR, 3′UTR, and A-tail (Fig. 4a). BA.5 mRNA vaccines were generated using pseudouridine triphosphate (Ψ-UTP) or UTP and encapsulated within lipid nanoparticles (LNPs) containing an ionizable lipid to facilitate mRNA release from endosomes (Extended Data Fig. 5). The mRNA vaccines used in animal studies had diameters of 109–136 nm and a polydispersity index of 0.102–0.187 (Extended Data Fig. 5c,f,i). These vaccines solely expressed the noncleaved full-length S protein in HEK293T cells (Extended Data Fig. 5b,e,h). Notably, pseudouridine-incorporated mRNAs exhibited slightly increased engineered S protein expression.

**Fig. 4.**
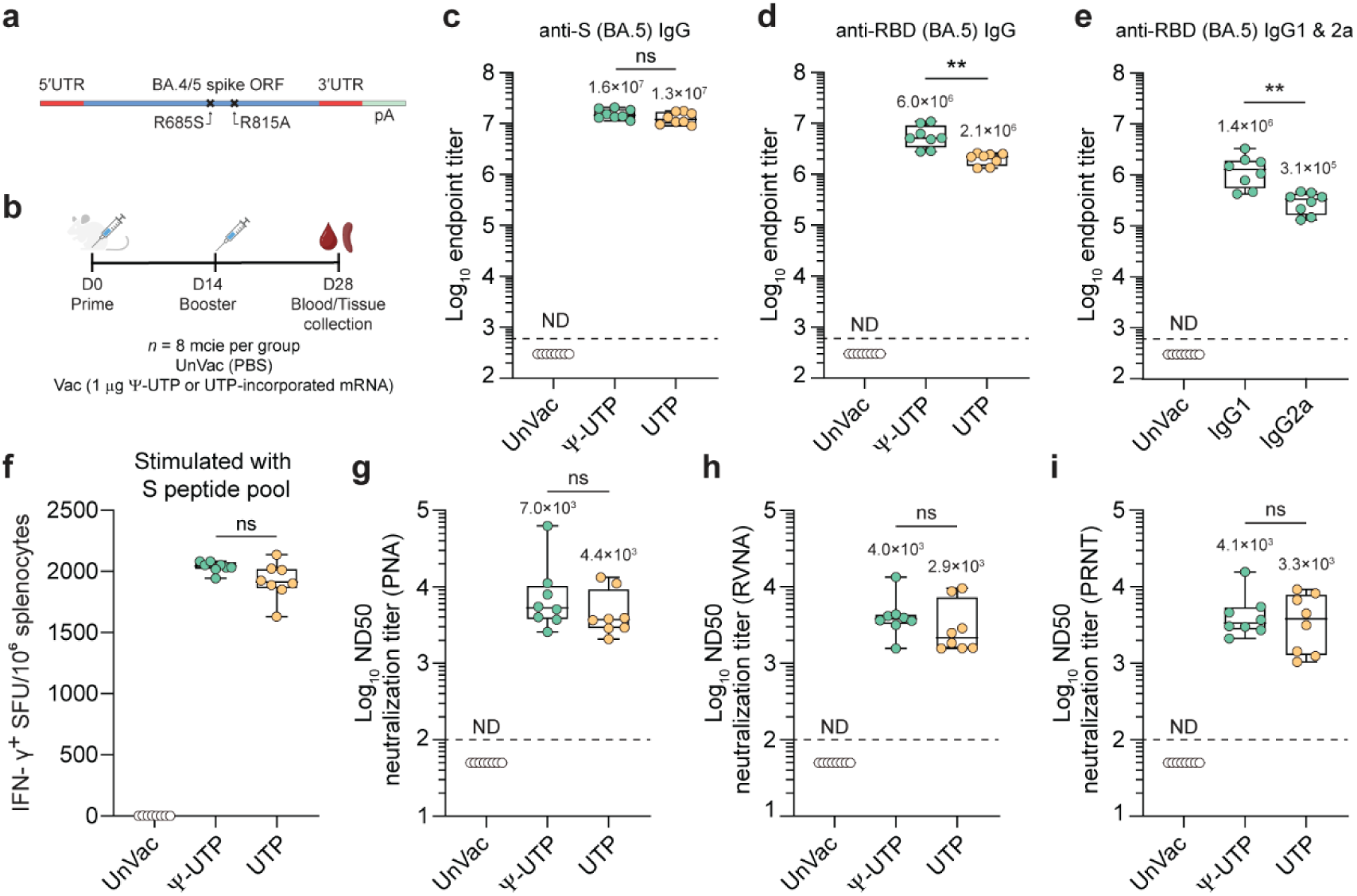
Immunogenicity of the noncleaved, nonfusogenic BA.5 S antigen with the SA substitution. **a**, Schematic representation of the mRNA used. **b**, Immunization and sample collection timeline. Seven-to-eight-week-old female BALB/c mice were immunized twice over a two-week interval with PBS (UnVac) or 1 μg of mRNA synthesized either with UTP or pseudouridine triphosphate (Ψ-UTP), encapsulated using an LNP (*n* = 8 per group). Two weeks after the final booster dose, mice were sacrificed, and sera and spleens were harvested. **c**,**d**, Levels of binding IgGs against Omicron BA.5 S (**c**) and RBD (**d**) in sera collected two weeks post booster immunization. **e**, Levels of Omicron BA.5 RBD-binding IgG1 and IgG2a in the sera from mice immunized with Ψ-UTP incorporated mRNA. **f**, Number of IFN-γ-producing cells per million splenocytes two weeks after the booster immunization, as measured by ELISpot. **g**–**i**, Neutralizing antibody levels (ND50) in mouse sera were quantified using three different neutralization assays with pseudovirus (**g**), reporter virus (**h**), and live virus (**i**). Each data point represents the calculated ND50 from biological triplicates for PNA, and biological duplicates for RVNA and PRNT. Data are represented as box-whisker plots (median, horizontal line; 25%–75% range, box; min-max values, whiskers), with individual data points superimposed on the graph. Statistical significance was determined by an unpaired Student *t*-test on the log_10_-transformed data. \*\**P* < 0.01; ns, not significant. The dotted lines indicate the assay limit of detection (LOD). Non-detected (ND) antibody levels are assigned values half of the LOD for graphical representation.

Both Ψ-UTP and UTP mRNA, at 0.5 μg dose in HEK293T cells and 0.25 and 1 μg doses in BALB/c mice, did not significantly induce inflammatory immune responses. However, the administration of 10 μg mRNA to mice via intramuscular injection led to the induction of proinflammatory cytokines IL-6, IFN-α, and IFN-γ, while having no impact on IL-1β and TNF-α levels (Extended Data Fig. 6).

Following two immunizations with a 2-week interval, Ψ-UTP- and UTP-incorporated mRNA vaccines at a dose of 1 μg elicited robust humoral immune responses in BALB/c mice (Fig. 4b). BA.5 S-binding IgG titers (reciprocal endpoint titer) exceeded 10^7^ for both mRNA vaccines (Fig. 4c), with higher RBD-binding IgG titers for Ψ-UTP-mRNA vaccine (2.9-fold, *P* = 0.0027) (Fig. 4d). We observed a balanced induction of high titer RBD-binding IgG1 and IgG2a, serving as surrogates for T_H_2- and T_H_1-mediated immune responses, respectively (Fig. 4e). ELISpot assays exhibited around 2000 SFUs per million splenocytes for both mRNA vaccines, suggesting robust S protein-specific T-cell responses. Anti-SARS-CoV-2 BA.5 neutralizing antibody (nAb) levels exceeded 10^3^ GMT, and no significant differences were observed in nAb levels between Ψ-UTP-mRNA and UTP-mRNA, as determined by the three different neutralization assays: pseudovirus neutralization assays (PNA), reporter virus neutralization assay (RVNA), and plaque reduction neutralization test (PRNT) (Fig. 4g–I, Extended Data Fig. 7).

### Protective immunity induced by BA.5 S(SA) antigen-encoding mRNA vaccines

Subsequently, we evaluated the protective immunity of the mRNA vaccine in 7-week-old BALB/c mice immunized with 1 μg mRNA at a two-week interval (Fig. 5a). After confirming high anti-RBD IgG levels (∼10^6^ reciprocal endpoint titers) induced in both mouse groups vaccinated with Ψ-UTP- or UTP-incorporated mRNA (Fig. 5b), these mice were intranasally challenged with 10^5^ PFU of Omicron BA.5 (NCCP 43426) on day 33. The BA.5 S protein carries the N501Y substitution, enabling the S variant to bind mouse ACE2^24^, rendering BALB/c mice susceptible to the Omicron BA.5 strain. As shown in Fig. 5c,d, lung tissue viral RNA copies significantly decreased in immunized groups compared to sham-vaccinated mice, with infectious virus titers below the detection limit of the plaque assay. On the second day post-infection, TNF-α and IFN-γ remained undetectable in both control and immunized groups, while serum IL-6 titers, an early cytokine response to viral infection^25^, notably rose in lung tissues of non-vaccinated mice (Fig. 5e–g). Moreover, lung sections stained with hematoxylin and eosin (H&E) exhibited alveolar damage near infected regions, accompanied by CD3-positive T-cell recruitment in vaccinated mice (Fig. 5h), demonstrating rapid memory T-cell activation in immunized mice. Collectively, these results demonstrate that BA.5 S(SA) mRNA vaccines synthesized with natural UTP or Ψ-UTP effectively establish protective immunity upon booster immunization.

**Fig. 5.**
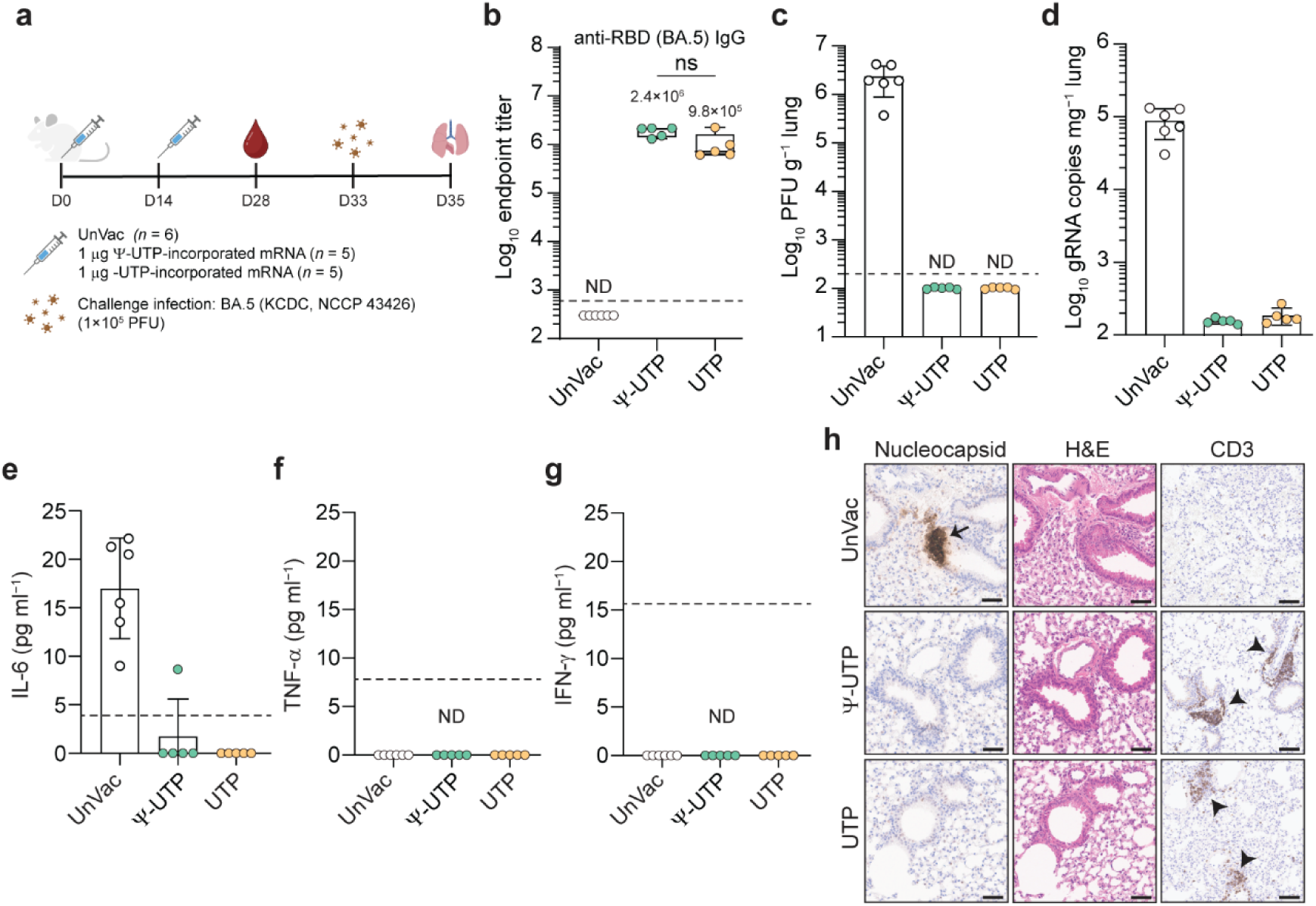
Protective efficacy of BA.5 mRNA vaccines expressing the S(R685S/R815A) antigen (BA.5_SA) tested in a BALB/c mouse challenge model. **a**, Schematic of the challenge experiment schedule. Eight-week-old female BALB/c mice were immunized as described in Fig. 4a (*n* = 5 for each group and *n* = 6 for UnVac). Two weeks after the booster dose, blood was collected via tail clip bleeding on day 28. Immunized mice were challenged intranasally with 1 × 10^5^ PFU of SARS-CoV-2 BA.5 on day 33. Mice were sacrificed 2 days post-infection to measure the tissue viral load in the lungs. **b**, Levels of binding IgG against Omicron BA.5 RBD in sera collected two weeks post booster immunization. **c**,**d**, Tissue viral load measured by plaque assay (**c**) and RT-qPCR (**d**) in the lung homogenate (tops of boxes indicate mean values). Each data point represents the viral load from two technical duplicates for RT-qPCR and two biological duplicates for plaque assay. In **b,c,** Non-detected (ND) antibody and viral load levels are assigned values half of the LOD for graphical representation. **e–g**, Cytokine levels in the lung homogenates were measured by ELISA. Each data point represents the cytokine concentration from two technical replicates. Samples with concentrations below the assay LOD were assigned a value of zero. **h**, Immunohistochemistry (IHC) and hematoxylin and eosin (H&E) staining of the lung sections. Lung sections for IHC were stained with a SARS-CoV-2 N-specific monoclonal antibody (tailed arrow) and CD3-specific polyclonal antibody (arrowheads). Scale bar, 100 μm. Statistical significance was determined by a Student *t*-test on the log_10_-transformed data. ns, not significant. The dotted lines indicate the LOD.

### Broad neutralizing efficiency of monovalent BA.5 S(SA) vaccines against various Omicron subvariants

The Omicron variant (B.1.1.529) and its descendants, carrying numerous mutations in the S protein, primarily concentrated in the RBD (Fig. 6a), have raised concerns due to their ability to evade the pre-existing immunity established against the ancestral S protein antigens^1,2,4,26–29^. We assessed the cross-reactivity of BA.5 S(SA) mRNA vaccines against various Omicron subvariants using S variant-pseudotyped MLVs (Extended Data Fig. 8). Both UTP- and Ψ- UTP-incorporated S(SA) mRNA vaccines showed reduced neutralizing effectiveness against BA.2.75.2 and XBB.1 variants, but they exhibited comparable efficacy against the BQ.1 variant harboring only two additional mutations (K444T and N460K) within the BA.5 RBD (Fig. 6b). BA.2.75.2, a variant of BA.2.75 (dubbed Centaurus), and the XBB.1 S variant, bearing 6 and 10 additional mutations within the immunodominant RBD, displayed a 3- and 6.2-fold reduction in ND50 levels (\*\*\**P* < 0.001 and \*\*\*\**P* < 0.0001) (Fig. 6b). The vaccines exhibited a similar attenuated neutralizing efficacy against the more recent XBB subvariants, XBB.1.5 and XBB.1.6 (Fig. 6c), with GMT ND50 for these three XBB variants yet exceeded 500.

**Fig. 6.**
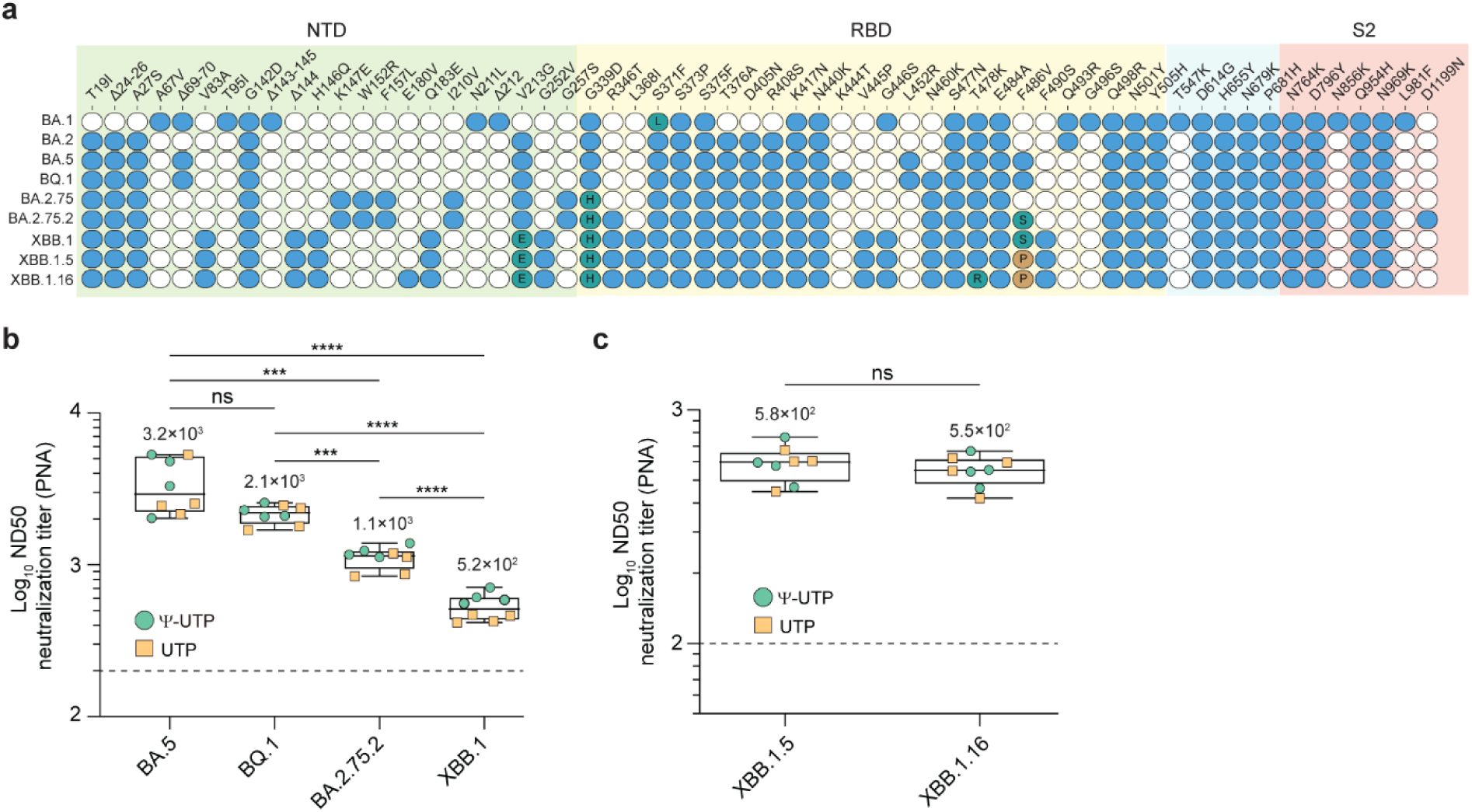
Broad neutralizing activity of monovalent BA.5_SA mRNA vaccines. **a**, Profile of amino acid mutations in the S protein of Omicron subvariants. **b**,**c**, Neutralizing antibody responses against Omicron subvariants. Neutralizing antibody levels (ND50) in sera collected from immunized mice (as described in Fig. 4b) were determined using pseudovirus neutralization assays. The box-whisker plots represent the median as a horizontal line and the 25th and 75th percentiles as the lower and upper borders of the box, with individual data points superimposed on the graphs along with the whiskers. Each data point represents the ND50 from three biological replicates. Statistical significance was assessed using one-way ANOVA on the log_10_-transformed data with Tukey’s multiple comparisons test. \*\*\**P* < 0.001; \*\*\*\**P* < 0.0001; ns, not significant. The dotted lines represent assay LOD. Non-detected (ND) antibody levels are assigned values half of the LOD for graphical representation.

## Discussion

In the present study, we demonstrated the immunogenicity of an Omicron BA.5 S(SA)-coding mRNA vaccine produced with ψ-UTP or UTP, providing protective immunity against BA.5. The R685S substitution incorporated into this vaccine antigen was found to be defective in furin-mediated pre-priming of SARS-CoV-2 S protein and showed an enhanced ACE2-dependent viral entry feature, along with reduced cell-to-cell fusion activity. The enhanced endosomal pathway-mediated viral entry in HEK293T cells expressing lower levels of ACE2 suggests an increased binding affinity of the S(R685S) variant to the ACE2 receptor, as previously observed with an S protein carrying a PRAR furin cleavage site deletion^30^. We found that the BA.5 S(S685S) variant, which still has the S2ʹ (R815) site remaining, displayed a substantially reduced fusogenic activity. Since the furin-mediated processing occurs in the ER-Golgi network, the reduced fusogenic activity caused by this mutation might, in part, be attributed to the change in the post-translational trafficking of the S to the cell surface^31,32^. Strikingly, the sensitive DSP-based fusion assay revealed that the S mutant with an R815A substitution also had a marginal, yet detectable, cell-cell fusion activity in HEK293T/hACE2 cells, consistent with previous findings^33^. Importantly, the R815A mutant we introduced in the S(SA) antigen was defective in generating the S2 fragment. The residual fusogenic activity of the R815A mutant was more evident when both ACE2 and TMPRSS2 were expressed abundantly in the acceptor cells. On the other hand, the S(SA) antigen did not induce any cell-cell fusion even under the same conditions optimal for inducing cell-cell fusion. The BA.4/5 S(SA) antigen, with an additional R815A mutation in combination with S(R685S) mutation, proved to be immunogenic, eliciting high titers of nAbs against multiple Omicron subvariants including the currently circulating XBB lineage variants.

Extracellular shedding of the cleaved S1 domain, caused by furin-mediated proteolytical cleavage of S protein can trigger systemic inflammatory responses when the S1 subunit dissociated from the S protein is bound by the TLR4 expressed on innate immune cells^19,20^. Thus, the S(SA) antigen may lower the risk of activating undesirable inflammatory responses. However, it is worth noting that the secreted S1 fragment bearing the RBD may enhance MHC II-mediated B-cell immune responses, which can be the case with the S(2P) antigen used in authorized mRNA vaccines^17^.

mRNA-LNP vaccines can activate the IFN-signaling pathways through the cytosolic foreign RNA sensors, RIG-I, MDA-5, and/or PKR, which can interfere with antigen expression^34^. Indeed, the two currently approved mRNA vaccines, mRNA-1273 and BNT162b2 vaccines have modified uridine incorporated in their mRNA in an attempt to evade the innate immune response and amplify antigen expression^16^. Our results showed that the 5′-capped mRNA vaccines prepared with natural UTP do not induce proinflammatory cytokine production *in vitro* (Extended Data Fig. 6) although S protein expression was slightly reduced (< 2-fold; Extended Data Fig. 5b,e,h). Furthermore, the UTP-incorporated mRNA vaccine did not induce proinflammatory cytokine production in mice receiving 0.25 or 1 μg mRNA. These results support the notion that natural UTP-incorporated mRNA itself, similar to cellular mRNAs, may pose little risk of inducing adverse innate immune responses as long as the mRNA is capped. It is also possible that the structural characteristics dictated by the sequence of the backbone and/or the antigen-coding region of the mRNA may influence the degree to which adverse inflammatory responses are triggered. Importantly, the incorporation of modified uridine nucleotides used to enhance antigen expression by stabilizing mRNA and possibly by suppressing the activation of IFN responses may introduce a potential risk of ribonucleotide misincorporation during *in vitro* transcription^35^ and non-complementary codon usage during translation^36^. Therefore, the use of nonmodified nucleotides in mRNA may indeed be worth considering to achieve authentic antigen or therapeutic protein expression.

Over the last three years, numerous SARS-CoV-2 variants have emerged^3^. Particularly, the recent Omicron variants pose challenges to natural immunity generated by prior infection or vaccination with ancestral S proteins^26,28,29,37^. The mRNA vaccines encoding the Wuhan-Hu-1 S(2P) S protein, which was initially used in authorized COVID-19 vaccines, exhibit significantly reduced cross-reactivity against the Omicron variants^5,26^. Omicron subvariants, including BQ.1, BA.2.75.2, and XBB.1, possess specific amino acid substitutions that allow them to evade humoral immunity induced by the ancestral S protein^2,27,38^. These subvariants exhibited significantly reduced cross-reactivity, even when tested against the sera from individuals immunized with bivalent mRNA vaccines containing the BA.5 S-coding mRNA^38^. Aligning with these findings. the potency of the BA.5 S(SA) monovalent mRNA vaccine was reduced by approximately 6-fold against the most immune evasive XBB lineage variants, such as XBB.1, XBB.1.5, and XBB.1.16. Nevertheless, it displayed broad neutralizing activity, at least against these currently prevalent Omicron subvariants.

In conclusion, our study underscores that the design of the S(SA) antigen, which eliminates fusogenic activity and prevents S1 subunit shedding, would contribute to the ongoing efforts in developing effective and safety-enhanced vaccines to combat the challenges posed by emerging SARS-CoV-2 variants.

## Methods

### Plasmids, antivirals, and antibodies

The chemically synthesized human codon-optimized S protein-coding cDNAs, which were designed based on the publicly available protein sequence in the National Center for Biotechnology Information database (ca. Wuhan-Hu-1: YP_009724390.1), were obtained from Twist Bioscience (South San Francisco, CA, USA) or through Bionics Inc. (Seoul, S. Korea). The synthetic DNA fragments were PCR-amplified and cloned into pcDNA3.1 vector (Invitrogen, Waltham, MA, USA; cat. no. V79020) using the In-Fusion HD cloning kit (Takara Bio, Kusatsu, Shiga, Japan; cat. no. 638947). Mutations were introduced into cDNAs using the QuikChange II XL Site-Directed Mutagenesis kit (Agilent, Santa Clara, CA, USA; cat. no. 200521) to construct expression vectors for S variants with a single amino acid change. The expression vectors for the S variants with multiple amino acid changes or deletions were generated using the In-Fusion HD cloning kit (Takara Bio). The pcDNA3.1-hACE2 used to express human ACE2 (hACE2) protein was constructed by inserting the PCR-amplified hACE2 cDNA (NP_001358344.1) between the Nhe I and Xho I sites. The plasmid expressing hTMPRSS2 with an N-terminal HA epitope tag was purchased from Sino Biological Inc. (Beijing, China; cat. no. HG13070-NY). The plasmids used for the dual split protein (DSP)-based cell-to-cell fusion assay were constructed as previously described^39,40^. Briefly, the *Renilla* luciferase (Rluc) gene was PCR-amplified from phRL-TK (Promega, Madison, WI, USA, cat. no. E6241), and the eGFP gene was PCR-amplified from a codon-optimized synthetic gene fragment obtained from Bionics Inc. (Seoul, S. Korea). The Rluc gene was split between the 229th and the 230th amino acid to make the two subunits, the N-terminal Rluc (nRluc) and the remaining C-terminal Rluc (cRluc). The GFP gene was split at the end of the 7th β sheet and the start of the 8th β sheet to make GFP1–7 (1–157 aa) and GFP8-11 (158–231 aa). pDSP1–7 was constructed by linking nRluc with GFP1–7 with a 4-amino acid residue linker (GLQG). pDSP8-11 was constructed by linking GFP8–11 and cRluc with a 2-aa residue linker (VD). Each gene fragment was PCR-amplified and cloned into a pcDNA 3.1 vector by using the In-Fusion HD cloning kit (Takara Bio). Lipofectamine 2000 (Invitrogen; cat. no. 11668019) was used for plasmid transfection.

Camostat (cat. no. SML0057) and E-64d (cat. no. E8640) were purchased from Sigma-Aldrich (St. Louis, MO, USA). A short synthetic dsRNA, poly(I:C) was obtained from Invivogen (San Diego, CA, USA; cat. no. TLRL-PICW).

Antibodies were obtained as follows: mouse monoclonal antibody targeting the C-terminal region of the SARS-CoV-2 S protein (GeneTex, Irvine, CA, USA; cat. no. GTX632604); rabbit polyclonal antibody targeting the N-terminal region of the SARS-CoV-2 S protein (GeneTex; cat. no. GTX135384); mouse monoclonal antibody against SARS-CoV nucleocapsid (N) protein (Sino Biological; cat. no. 40143-MM08); rabbit polyclonal antibody targeting N-terminus region of human ACE2 (Genetex; cat. no. GTX101395); rabbit polyclonal antibody against human GAPDH (Abcam, Cambridge, UK; cat. no. ab9485); mouse monoclonal antibody targeting the 872–891 amino acid sequence within the human α-actinin (Santa Cruz Biotechnology Inc, Dallas, TX, USA; cat. no. sc-166524); mouse monoclonal antibody against MLV p30 (Abcam; cat. no. ab130757).

### Cell culture and virus infection

HEK293T (human embryonic kidney-derived cells), Calu-3 (human lung adenocarcinoma cells), A549 (adenocarcinomic human alveolar basal epithelial cells), and VeroE6 (African green monkey kidney cells) cell lines were obtained from the American Type Culture Collection (ATCC, Rockville, MD, USA). The VeroE6 cell line stably expressing hTMPRSS2 (Vero E6/TMPRSS2) was a gift from Dr. Seong-Jun Kim (Korea Research Institute of Chemical Technology, KRICT, Daejeon Korea). The VeroE6 cell line stably expressing hTMPRSS2 and hACE2 (Vero E6-TMPRSS2-T2A-ACE, referred to as VeroE6/A2-T2) was obtained through BEI Resources, NIAID, NIH (cat. no. NR-54970). The HEK293T/hACE2 and A549/hACE2 cell lines, which stably express human ACE2 receptor, were established by transducing HEK293T and A549 cells with an hACE2-expressing recombinant lentivirus and selecting a single cell clone by limiting dilution of the transduced cells in the presence of 2 or 3 μg ml^−1^ puromycin. Third-generation lentiviral transfer vector encoding hACE2, the second-generation lentiviral packaging plasmid (Addgene plasmid #12260), and a VSV G-expressing vector (Addgene plasmid #12259) were used to prepare the lentiviruses used in transduction experiments.

The HEK293T, Vero E6, Calu-3, and A549 cells were cultivated in Dulbecco’s modified Eagle’s medium (DMEM) supplemented with 10% fetal bovine serum (FBS), 100 U ml^−1^ of penicillin, and 100 μg ml^−1^ streptomycin at 37 °C in 5% CO_2_. The A549/hACE2 cells stably expressing human ACE2 receptors were maintained in a complete medium with 3 µg ml^−1^ of puromycin. Vero E6/TMPRSS2 and VeroE6/A2-T2cells were maintained in a complete medium with 150 mg ml^−1^ of hygromycin and 10 µg ml^−1^ of puromycin, respectively.

All SARS-CoV-2 viruses used in this study (two S-clade strains, NCCP 43326 and NCCP 43331; Omicron BA.5 strain, NCCP 43426) were obtained from the National Culture Collection for Pathogens, Korean Center for Disease Control (Osong, S. Korea). SARS-CoV-2 was propagated in VeroE6 cells for S-clade and VeroE6/TMPRSS2 cells for Omicron BA.5. Cells were infected at a multiplicity of infection (MOI) of 0.01. Virus titer was determined by plaque assay on VeroE6 cells for S-clade SARS-CoV-2 and VeroE6/TMPRSS2 cells for Omicron BA.5 as previously described^41^.

### S protein structure modeling

AlphaFold2, provided by Neurosnap Inc., was used to predict the monomeric structure of the S protein^42^, with the highest-ranking model selected as the representative image.

### RNA sequencing analysis

Viral RNA from the original P0 (10 μl) or P1 sample (125 μl) was extracted using TRIzol LS reagent (Invitrogen; cat. no. 10296028). RNA was subjected to variant calling sequencing using barcode-tagged contiguous sequencing (BTseq) by Celemics Inc., Seoul, Korea.

### Real-time reverse-transcription quantitative PCR (RT-qPCR)

Total RNA from cells or tissue homogenate was extracted using TRIzol reagent (Invitrogen; cat. no. 15596026). The SARS-CoV-2 genomic RNA copy number was determined by RT-qPCR using a specific set of primers and a probe targeting the ORF1ab along with the real-time PCR Master Mix (TOYOBO, Osaka, Japan; cat. no. QPK-101) as described previously^41^. Pseudovirus genome copy number was quantified using RT-qPCR. Packaged viral RNA was extracted using TRIzol LS reagent (Invitrogen) and subjected to RT-qPCR using primers targeting the 5′ LTR of MLV RNA^43^. Standard RNAs were prepared through *in vitro* transcription using a T7 MEGAscript kit (Invitrogen; cat. no. AM1334), followed by electrophoresis on an 8 M urea-5% polyacrylamide gel and purification via gel-extraction.

TNF-α and IFN-β mRNA expression levels were determined using the ΔΔC_t_ method and were normalized to the expression level of GAPDH mRNA, as described^44^.

### Immunoblotting

Cells were lysed with RIPA buffer (150 mM NaCl, 1.0% NP-40, 0.5% sodium deoxycholate, 0.1% SDS, 50 mM Tris, pH 8,0, 1× protease inhibitor cocktail) and centrifuged at 13,000*g* for 15 min at 4 ℃. For the analysis of S proteins loaded onto pseudoviruses, the MLV-containing culture supernatant was subjected to ultracentrifugation (150,000*g* for 2 h at 4 ℃) over 20% (w/v) sucrose cushion in a TL-100 centrifuge using the TLA100.3 rotor to collect pseudovirus pellets, as described^45^. Subsequently, both pseudovirus samples and cell lysates were subjected to SDS-PAGE, and the proteins were transferred to a PVDF membrane (GE Healthcare Life Sciences, Chicago, IL, USA; cat. no. 10600023). The membrane was then blocked with 5% bovine serum albumin and probed with the appropriate set of primary and secondary antibodies.

### Pseudovirus entry assay

SARS-CoV-2 S protein (with a deletion of the C-terminal 19-amino acids ER-retention signal)-pseudotyped murine leukemia virus (MLV) was produced as previously described^41,46^.

### Cell-to-cell fusion assay

The DSP-based SARS-CoV-2 S fusogenic activity assay was performed as previously described^47^. Briefly, HEK293T cells co-transfected with pDSP1–7 and a full-length S protein-expression vector (donor cells) and HEK293T cells expressing hACE2 and pDSP8–11 (acceptor cells) were co-cultured in a 12-well plate with a total seeding density of 1.5 × 10^6^, by mixing the donor and acceptor cells at a 1:1 ratio at 24 h post-transfection. After 12 or 24 h, the cells were harvested and subjected to a luciferase assay using the Rluc Assay System (Promega; cat. no. E2820). Luminance (ALU) measured in a GloMax Navigator Microplate Luminometer (Promega) was normalized to microgram lysate protein.

Brightfield microscopy was employed to visualize the fusion-induced multicellular syncytia using an inverted phase contrast microscope (OPTIKA, Ponteranica, Italy). Confocal microscopy was performed to quantity the reconstituted GFP fluorescence generated by fusion between donor and acceptor cells (1.8 × 10^5^ each per well) co-cultured in a collagen (Thermo Fisher Scientific; cat. no. A1048301) pre-coated (5 μg cm^−2^) 4-well chamber slide (Thermo Fisher Scientific, cat. no. 154526). A fluorescence image was captured using an LSM 880 confocal microscope (Carl Zeiss AG, Jena, Germany). Image reconstruction was conducted using the ZEN software (Zeiss).

### mRNA-LNP preparation

The BA.4/5 S-encoding plasmid, which has a template for 3′-end A-tailing (approximately 120-nt), was linearized with BbsI, purified using the Gel Extraction kit (Bionics; cat. no. BNROP-0020), and used as a template for *in vitro* transcription with the MEGAscript T7 transcription kit. When indicated, ψ-UTP (TriLink, San Diego, CA, USA; cat. no. N-1019) was used to substitute for UTP to prepare ψ-UTP-incorporated mRNA. The CleanCap Reagent AG (3′ OMe) (TriLink; cat. no. N-7413) was added to the *in vitro* transcription reaction mixture for co-transcriptional 5′-end capping. The capped, A-tailed mRNAs were resolved on formaldehyde-denaturing agarose (0.7%) gel and stained with ethidium bromide to assess RNA integrity before use.

mRNA was transfected into cells using either Lipofectamine 2000 (Invitrogen) or LNPs. mRNA encapsulation within LNPs was carried out using the NanoAssemblr Spark (Precision Nanosystems, Vancouver, Canada). Briefly, a mixture of ionizable (ALC-0315; MedChemExpress, cat no. HY-138170), structural (cholesterol, Sigma-Aldrich; cat. no. C8667), and helper [distearoylphosphatidylcholine (DSPC), Avanti Polar Lipids, Inc., Alabaster, AL, USA; cat. no. 85030] lipids along with polyethylene glycol (PEG)-conjugated lipid (ALC-0159, MedChemExpress; cat. no. HY-138300) were combined with the mRNA in citrate buffer (pH 4) at a 1:6 ration (mRNA:lipid, molar ratio). The resulting encapsulated mRNA-LNP mixture was subjected to dialysis against PBS with 8% sucrose, as described^48^. mRNA encapsulation efficiency was calculated using the Quant-it RiboGreen RNA Assay kit (Invitrogen; cat. no. R11490) as described^49^. mRNA-LNP particle size and uniformity were evaluated by dynamic light scattering using a particle size analyzer (ELSZ-1000, Otsuka Electronics, Osaka, Japan).

### Cytokine ELISAs

Serum cytokine levels were detected via ELISA one day after mRNA-LNP immunization. To measure lung tissue cytokine levels in mice challenged with SATS-CoV-2, lung homogenates were incubated with Triton X-100 at a final concentration of 1% for 1 h at room temperature to inactivate the virus. The concentrations of mIFN-α, mIL-1β, mTNF-α, mIFN-γ, and mIL-6 in sera were determined using the ELISA kits obtained from PBL Assay Science (Piscataway, NJ, USA; cat. no. 42120-1 for mIFN-α), Elabscience (Houston, Texas, USA; cat. no. E-EL-M0037 for mIL-1β), and Thermo Fisher Scientific (cat. no. 88-7324-88 for mTNF-α, cat. no. 88-7314-88 for mIFN-γ, and cat. no. 88-7064-88 for mIL-6) according to the manufacturer’s recommendation.

### IgG ELISAs

SARS-CoV-2 S and RBD-specific IgGs were detected via ELISA. Plates were coated with 1 μg ml^−1^ of the antigen protein (Sino Biological; cat. no. 40589-V08H33 for BA.5 S and cat. no. 40592-V08H131 for BA.5 RBD) in phosphate buffer saline (PBS) overnight at 4 °C. Plates were then blocked for 6 h with PBST (PBS with 0.01% Tween-20) containing 2% BSA (Sigma-Aldrich; cat. no. 10735094001) at room temperature. After blocking, plates were washed with PBST and incubated with mouse sera diluted in the blocking buffer for 3 h at room temperature on a rocking stand. Following incubation with the sera, plates were washed four times with PBST, and α-S or α-RBD IgG bound to the plate was then detected using HRP-conjugated goat anti-mouse IgG (H+L) antibody (Invitrogen; cat. no. G21040), anti-mouse IgG1 antibody (Invitrogen; cat. no. A10661), or anti-mouse IgG2a antibody (Invitrogen; cat. no. M32207) at a 1:1500 dilution for 1.5 h at room temperature on a rocking stand. After washing the plates four times with PBST, 100 μl of TMB substrate (Thermo Fisher Scientific; cat. no. 34028) was added to each well for the detection of HRP-conjugated antibodies. The reaction was stopped by adding 1 M phosphoric acid, and the absorbance at 450 nm was read using the GloMax-Multi Detection System (Promega).

Endpoint titer was defined as the serum dilution fold that yielded an absorbance equal to the cut-off value. The cut-off optical density value was set to the plate background OD_450nm_ detected at the highest dilution of the sample sera plus 10× standard deviation, as previously described^15^. Endpoint titer was determined by plotting a dose-response curve (OD_450 nm_ vs. reciprocal serum-dilution fold) and four-parameter nonlinear regression analysis using the GraphPad prism 8·0 (GraphPad Software Inc., San Diego, CA, USA). The dose-response nonlinear regression curve equation and the equation used to determine S-binding IgG endpoint titer by extrapolation are as follows:

A nonlinear regression equation for the dose-response curve:

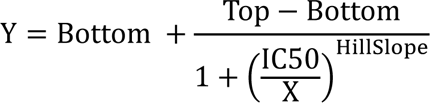

An equation for endpoint titer determination:

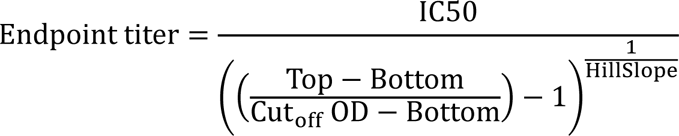

### Pseudovirus neutralization assay (PNA)

A neutralization assay using SARS-CoV-2 S protein-pseudotyped MLV (SARS2pp) was performed to determine neutralizing antibody titers in the sera, as previously described^46^. Briefly, equal volumes (60 μl each) of diluted mouse sera and diluted SARS2pp, which yields about 1 × 10^6^ ALU per well, were mixed (*n* = 3) and incubated for 1 h at 37 ℃. Then VeroE6/A2-T2 cells, which were grown for 24 h following seeding at 1.5 ×10^4^ per well on 96-well white plates, were transduced with the pseudotyped virus and mouse sera mix (100 μl per well). Following a media change at 12 h post-transduction, cells were further cultivated for 36 h and harvested for luciferase assay using the Nano-Glo luciferase assay system (Promega; cat. no. N1120) and the GloMax Navigator Microplate Luminometer (Promega). The 50% neutralizing dilution (ND50), which was defined as the serum dilution causing a 50% reduction in luciferase activity compared to the wells displaying no inhibition, was determined by plotting a dose-response curve (log_10_ ALU vs. reciprocal serum-dilution fold) and four-parameter nonlinear regression analysis using the GraphPad prism 8·0 (GraphPad Software Inc). The dose-response nonlinear regression curve equation and the rearranged equation used to determine the ND50 by extrapolation are as follows:

A nonlinear regression equation for the dose-response curve:

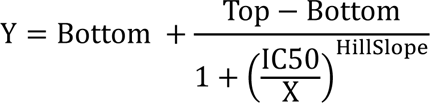

An equation for ND50 determination:

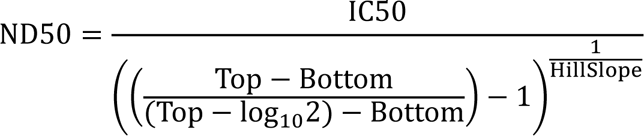

### Reporter virus neutralization assay (RVNA)

A nanoluciferase (Nluc)-expressing live, chimeric SARS-CoV-2 carrying the Omicron BA.5 *S* gene (derived from the BA.5 strain NCCP 43426) was generated using a full-length cDNA of the G-clade SARS-CoV-2 YS006^23^ that was cloned in a bacterial artificial chromosomal plasmid to obtain pBAC-SARS-CoV-2/YS006(BA.5_S)_Nluc, in which the ORF7 was replaced by the Nluc gene, as previously described^50^. The recombinant chimeric virus, rYS006(BA.5_S)_Nluc, was rescued by co-culturing of BHK-21 cells transfected with pBAC-SARS-CoV-2/YS006(BA.5_S)_Nluc with VeroE6/TMPRSS2 cells at 6 h post-transfection, as previously described^50^. The co-cultured cells were incubated until clear signs of cytopathic effects (CPE) were observed. Once significant CPE was observed in the coculture, supernatant (passage 0, P0 viral stock) was harvested and passaged onto VeroE6/TMPRSS2 cells to obtain the P1 viral stock used in the neutralization assay. Equal volumes (60 μl each) of diluted mouse sera and diluted rYS006(BA.5_S)_Nluc (95 PFU per well), which yields about 1 × 10^6^ ALU per well, were mixed (*n* = 3) and incubated for 1 h at 37 ℃. Then, the mixture (100 μl) was used to infect VeroE6/TMPRSS2 cells, which were grown for 24 h following seeding at 1.5 × 10^4^ per well on 96-well white plates. Growth media was changed 1 h post-infection and cells were further cultivated for 24 h. Harvested cells were subjected to luciferase assay. The ND50 was determined using the same method used in the PNA assay described earlier.

### Live virus plaque reduction neutralization (PRNT)

SARS-CoV-2 Omicron BA.5 variant (NCCP NCCP 43426) propagated in VeroE6/TMPRSS2 cells, which form ∼ 40 plaques in each well of a 12-well plate, was mixed with serially diluted mouse sera and incubated for 1 h at 37 °C. After incubating the mixture with VeroE6/TMPRSS2 cells for infection, the cells were overlaid with 1% agarose. When plaques became visible, cells were fixed with 10% formaldehyde and stained with 1% crystal violet. The ND50, which was defined as the serum dilution causing a 50% reduction in plaque number compared to the wells displaying no inhibition, was determined by four-parameter non-linear regression analysis of a dose-response curve (% neutralization vs. reciprocal serum-dilution fold) using the GraphPad prism 8·0 (GraphPad Software Inc). The % neutralization at each serum dilution = (plaque number formed w/o sera− plaque number formed w/ sera)/(plaque number formed w/o sera)

### Interferon-γ (IFN-γ) enzyme-linked immunosorbent spot (ELISpot) assay

ELISpot assay was carried out using murine IFN-γ single-color enzymatic ELISpot assay kit (ImmunoSpot, Cleveland, OH, USA) according to the manufacturer’s instruction. Briefly, spleens from immunized mice were harvested two weeks after the last vaccination. The splenocytes of immunized mice were then plated on IFN-γ antibody-coated PVDF-backed plates and stimulated with 0.1 μg ml^−1^ overlapping peptide pools spanning the full-length SARS-CoV-2 BA.4/5 S protein (JPT, Berlin, Germany; cat. no. PM-SARS2-SMUT-10). After overnight incubation at 37 °C in a humidified atmosphere containing 5% CO_2_, the plates were washed with PBS and PBST and the secreted IFN-γ was detected with a set of detection antibodies. The number of spot-forming units (SFUs) was counted automatically using an ELISpot reader (Immunospot Analyzer, ImmunoSpot).

### Animal experiments

All animal experiments were performed following the Korean Food and Drug Administration guidelines. Experimental procedures were reviewed and approved by the Institutional Animal Care and Use Committee of the Yonsei Laboratory Animal Research Center (Permit No: IACUC-A-202203-1427-02). At the termination of experiments, all mice were euthanized by CO_2_ inhalation.

BALB/c mice (7–8-week-old) were immunized by intramuscular injection of mRNA-LNP (50 μl) into the ipsilateral hind limb for prime-boost vaccination, using an insulin syringe (BD, Franklin Lakes, NJ, USA; cat. no. 328868). Blood samples were obtained through tail-clip bleeding or cardiopuncture. The collected blood samples were left to clot for 30 min at room temperature and were centrifuged at 6,000*g* for 25 min. Before use, all mouse sera were heat-inactivated by incubating for 30 min at 50 °C.

### Challenge infection

A vaccine efficacy study using BALB/c mice was carried out in a biosafety level 3 (BL3) facility at the Avison Biomedical Research Center (ABMRC), Yonsei University College of Medicine. The study was conducted under IBC permit number A-202009-260-01. The animal experiments involving SARS-CoV-2 were approved by the IACUC at Yonsei University College of Medicine (IACUC number: 2020-0227). Animals received two (D0 and D14) IM immunizations of mRNA-LNP vaccine or PBS. BALB/c mice were intranasally challenged with 1 ×10^5^ PFU of SARS-CoV-2 Omicron BA.5 variant. BALB/c mice were euthanized 2 days post-infection and lung viral titer was determined by performing plaque assay using VeroE6/A2-T2 cell and qRT-PCR of the lung homogenate. Four right lung lobes (superior, middle, inferior, and post-caval) were homogenized in serum-free DMEM using a bead beater. The resulting homogenate was clarified by centrifugation at 4 °C for 15 min. Supernatants were collected and subjected to viral titer analysis where viral RNA was detected via qRT-PCR and infectious viral load was determined via PRNT.

### Immunohistochemical analysis

The left lung lobe was subjected to histological analysis. Paraffin blocks containing the left lung lobe were sectioned at approximately 4 microns and stained using hematoxylin and eosin (H&E) or immunostained using a rabbit monoclonal antibody against SARS-CoV-2 nucleocapsid protein (Sino Biological; cat. no. 40588-R002) and rabbit polyclonal antibodies against CD-3 protein (Dako, Glostrup, Denmark; cat. no. A0452).

### Statistical analysis

Results are presented as the mean ± standard deviation (SD). Statistical analyses were performed using GraphPad Prism 8 (GraphPad Software Inc.). Differences between groups were considered statistically significant at *P* < 0.05. (\**P* < 0.05; \*\**P*< 0.01; \*\*\**P* < 0.001; \*\*\*\**P* < 0.0001; ns, not significant.)

## Material availability

All reagents generated in this study will be made available on request after completion of a Materials Transfer Agreement.

## Data availability

Source data and images are provided with this paper. The authors declare that any additional information supporting the findings of this study is available from the corresponding author upon reasonable request.

## Code availability

No code was used in the course of the data acquisition or analysis.

## Acknowledgments

We thank the National Culture Collection for Pathogens (NCCP) at the Korean Center for Disease Control (Osong, S. Korea) for providing SARS-CoV-2 viral stocks. This work was supported by the National Research Foundation of Korea (NRF) grants (2020R1A2C2005170 and 2022M3E5F1016361) funded by the Ministry of Science and ICT (MIST), South Korea. H.C. was partially supported by a postdoctoral fellowship from the Brain Korea 21 (BK21) FOUR program.

## Author contributions

H.Y.S., and J.-W.O. conceived and designed the experiments. H.Y.S., H.J., H.W., H-G.J., H.C., Y.B., S-Y.L., G.Y., and S-Y.H. performed experiments. H.Y.S, Y-M.S. and H-G.J. prepared the mRNA-LNP vaccine. H.Y.S., and J.-W.O. analyzed the data. H.Y.S., H.J., and Y.B. carried out the infection experiments with SARS-CoV-2 conducted at the BSL3 facility. I.P., and J.-S.S. supervised the experiments carried out in the BL3 facility. H.Y.S., and J.-W.O. wrote the paper with input from other authors. J.-W.O. supervised the studies.

