## Supplementary Information for "Enhanced Omicron subvariant cross-neutralization efficacy of a monovalent SARS-CoV-2 BA.4/5 mRNA vaccine encoding a noncleaved, nonfusogenic spike antigen"

**Extended Data Figures** 1–8
**Supplementary Tables** 1 and 2

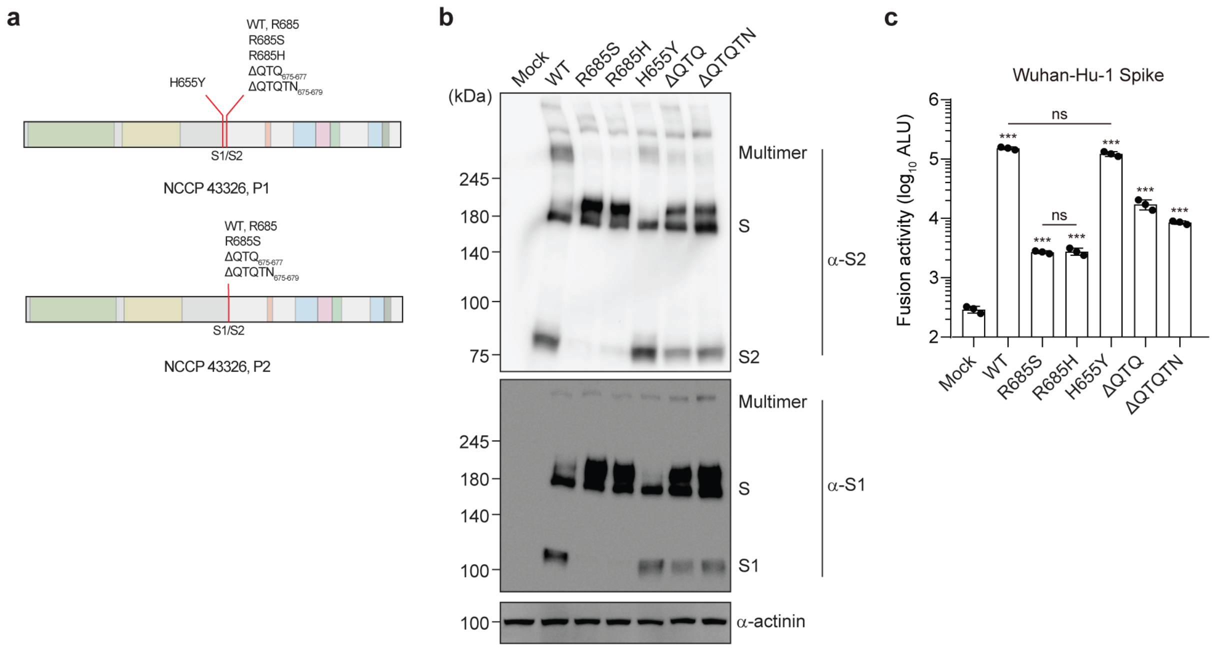


**Extended Data Fig. 1. Profile of S variants in the VeroE6-passaged NCCP43326 strain. a**, S variants in the P1 and P2 stocks of the S clade SARS-CoV-2 isolate NCCP43326. **b**, Profile of S1/S2 cleavage efficiency of the representative S variants identified through RNA sequencing. **c**, Cell-cell fusion activity of the representative S variants. The DSP-based fusion assay was conducted as described in Fig. 2d.

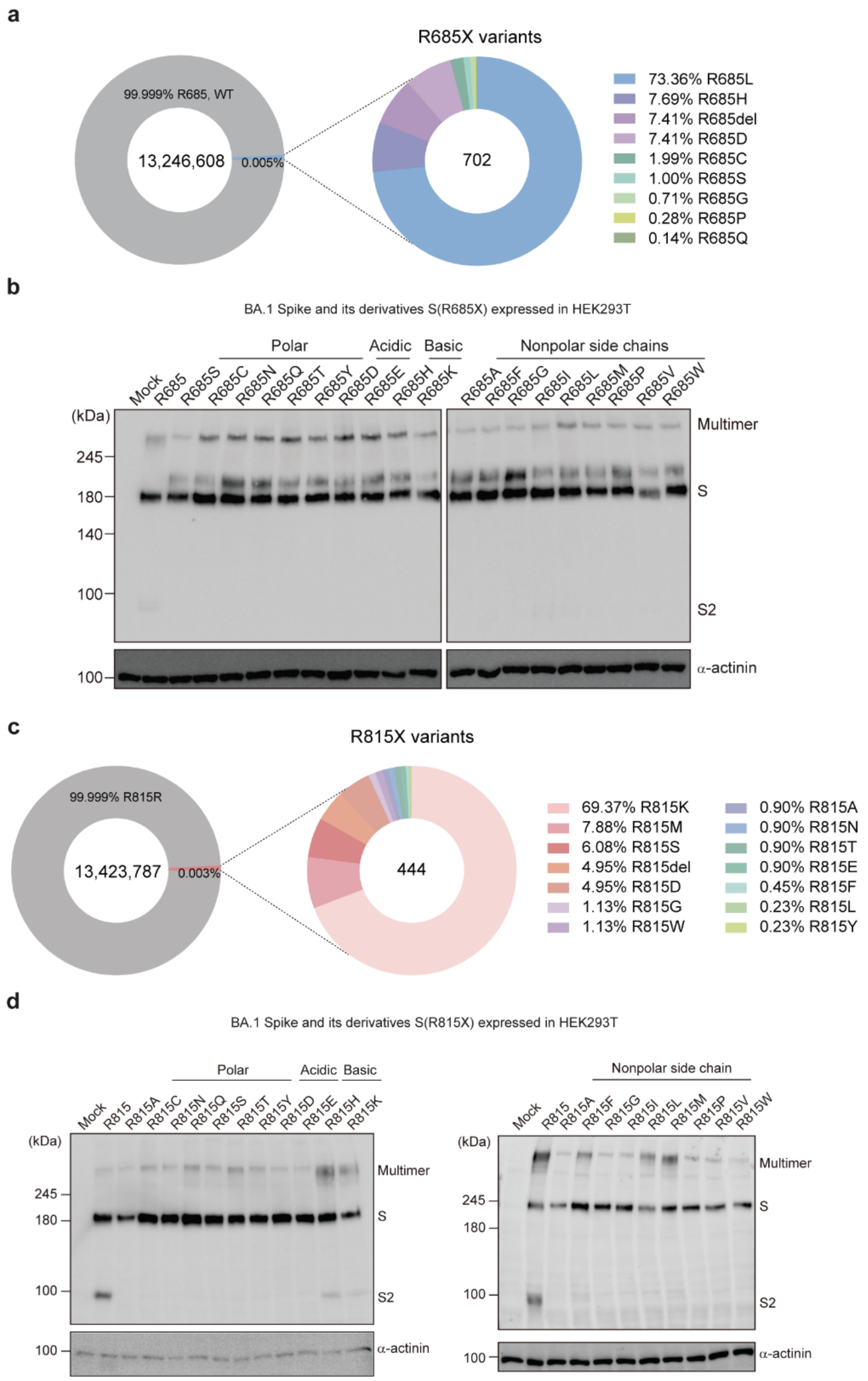


**Extended Data Fig. 2. Effect of R685 and R815 mutations on Omicron BA.4/5 spike protein cleavage in HEK293T cells. a,c**, Profile of amino acid variations at the R685 residue (**a**) or the R815 site (**c**) in S protein variants. Donut charts displaying R685X or R815X mutations in SARS-CoV-2 genome sequence data sets from the GISAID database (as of February 1, 2023). **b**,**d**, Western blot analysis of HEK293T cells expressing ectopically the indicated S variants for 2 days. The SARS-CoV-2 S protein was detected using antibodies specific to the C-terminal region of the protein.

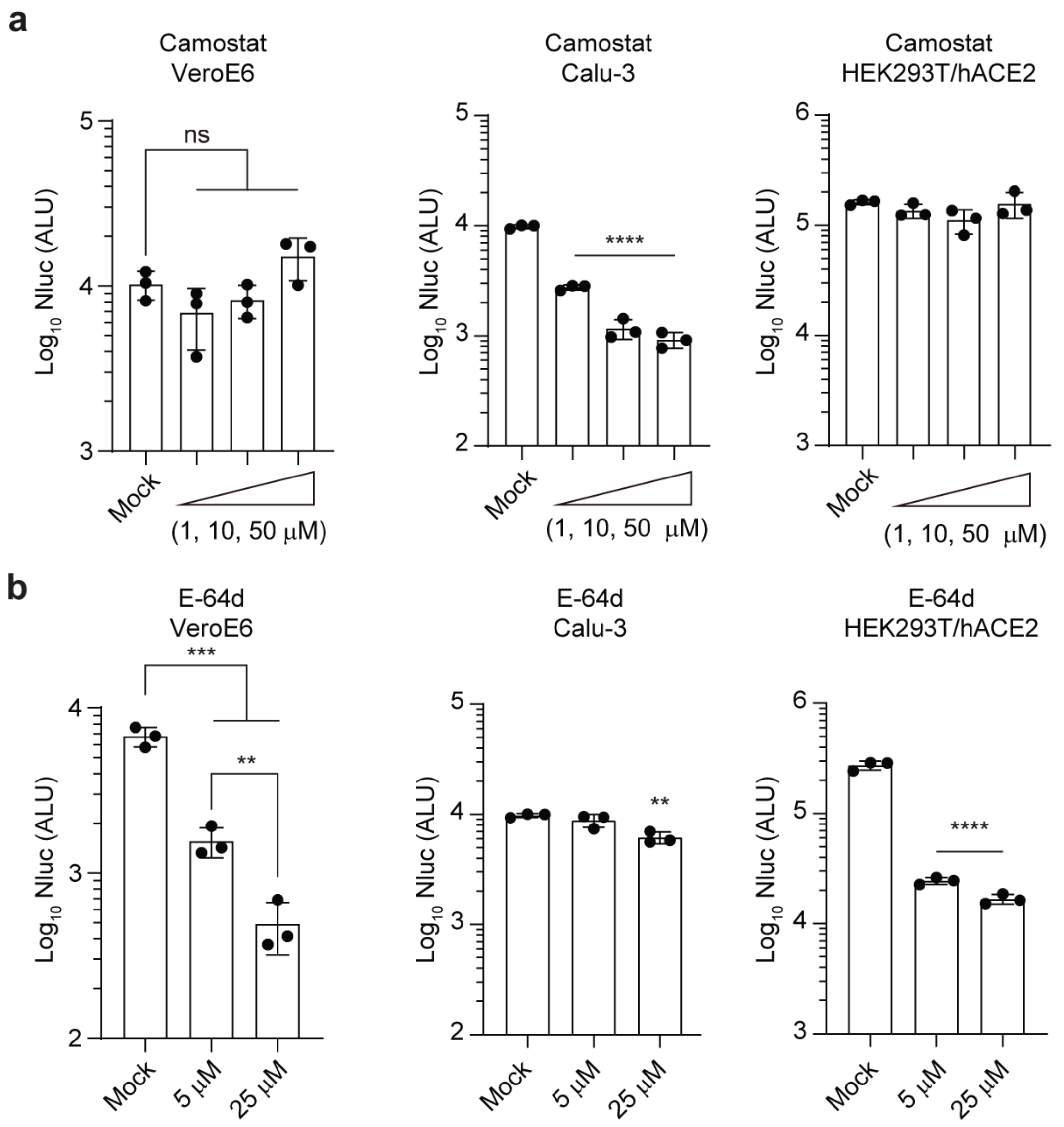


**Extended Data Fig. 3. Differential inhibitory activity of camostat and E-64d in various cell types.** Wuhan-Hu-1 S protein-pseudotyped MLV was transduced into the indicated cell lines. After 24 h of drug treatment, nanoluciferase activity was assessed. Each data point represents values from biological triplicates. Statistical significance was determined using an unpaired Student *t*-test. ** $P$ <0.01; *** $P$ < 0.001; **** $P$<0.0001; ns, not significant.

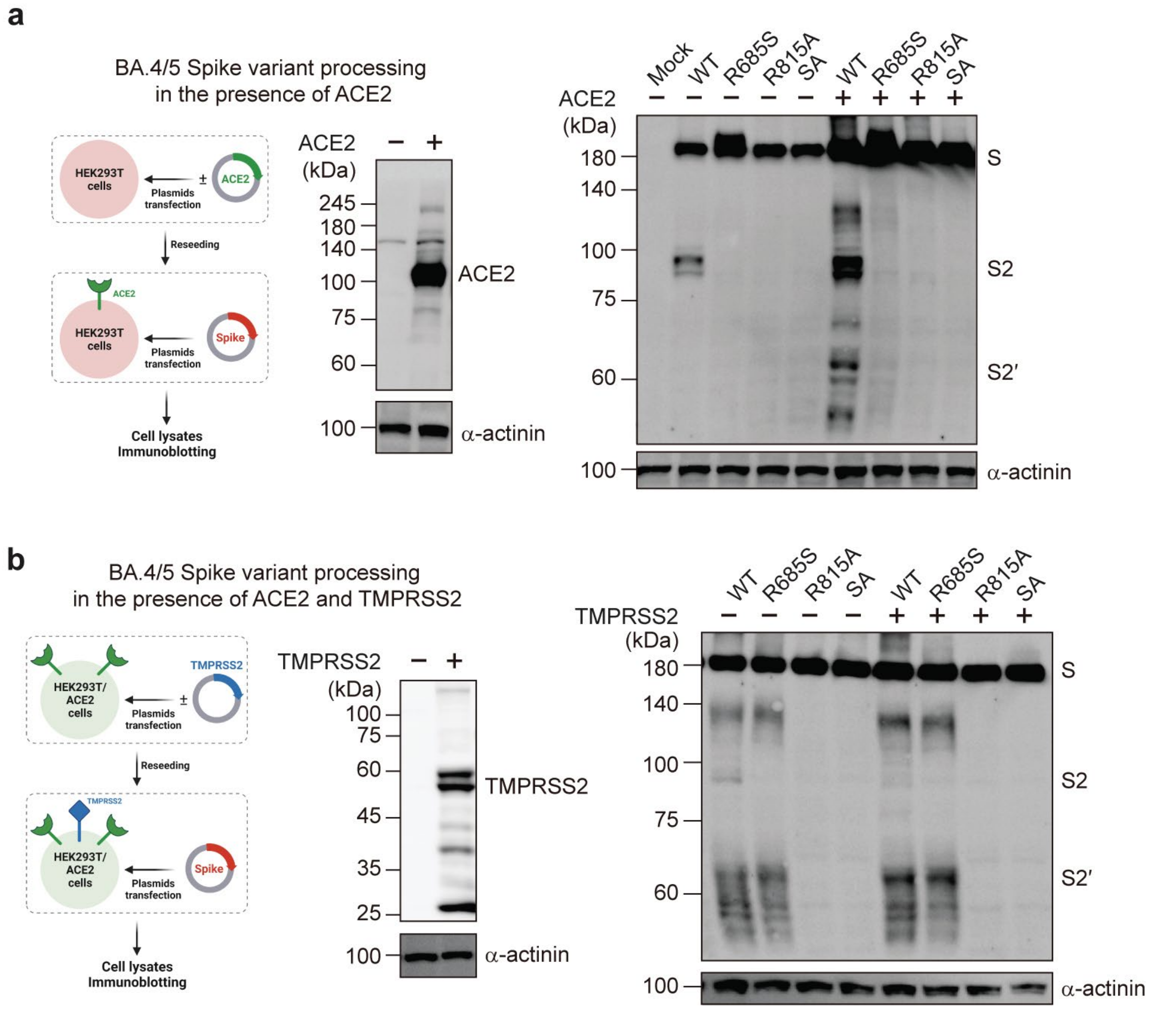


**Extended Data Fig. 4. Effect of ACE2 and TMPRSS2 on the processing of Omicron BA.4/5 S protein and its variant at the S1/S2 and S2′ cleavage sites.** HEK293T cells were transfected with or without an hACE2-expressing vector (**a**), or HEK293T/hACE2 cells were transfected with or without a TMPRSS2-expressing vector (**b**). These cells reseeded in 6-well plates were further transfected with a vector expressing the WT BA.4/5 S or its variants with single mutations (R685S or R815A) or double mutations (R685S and R815A, SA) at 12 h post-transfection. Two days later, cells were subjected to Western blot analysis. SARS-CoV-2 S protein was detected using antibodies specific to the C-terminal region of the protein, and α-actinin was utilized as a loading control to ensure equal protein loading across samples. The experimental schemes were created with BioRender.com.

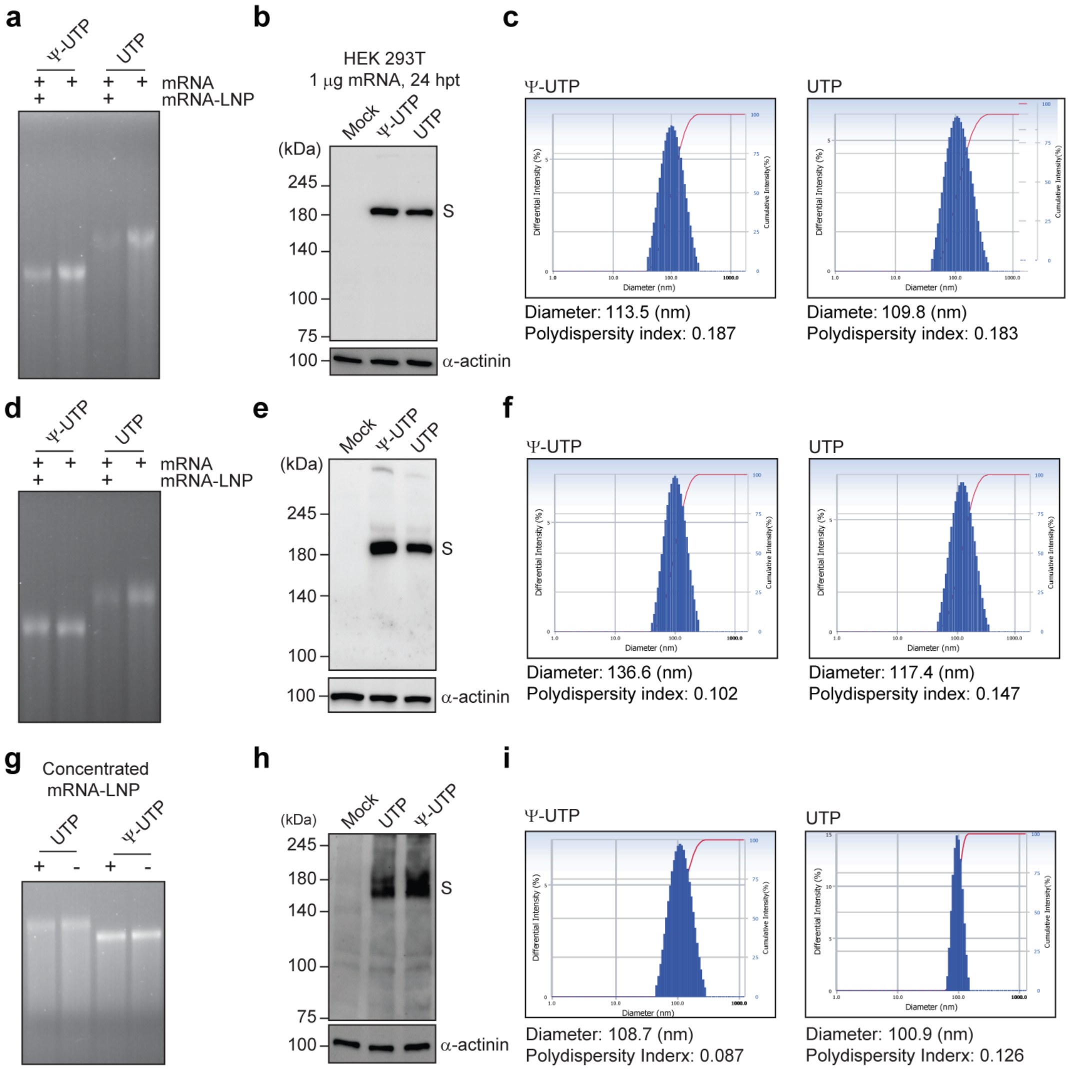


**Extended Data Fig. 5. Features of the BA.4/5 S(SA) mRNA-LNP vaccines used in immunization.** The mRNAs encoding the BA.4/5 spike SA antigen, prepared using UTP or pseudouridine triphosphate ψ-UTP, were then formulated with an ethanol-lipid mix. **a**–**c**, The mRNA-LNPs used for prime immunization. **d**–**f**, The mRNA-LNPs used for booster immunization. **g**–**i**, The concentrated mRNA-LNPs used to evaluate their innate immune activation capacity. **a**,**d**,**g**, Integrity of the mRNAs before and after LNP formulation, visualized by ethidium bromide staining following electrophoresis on a denaturing formaldehyde agarose gel. **b**,**e**,**h** Expression of the spike antigen in HEK293T at 24 h post-transfection of the formulated mRNA (1 μg)-LNP, analyzed by immunoblotting. α-actinin served as a loading control. **c**,**f**,**i**, Particle size and uniformity of the mRNA-LNPs, analyzed using a dynamic light scattering particle size analyzer.

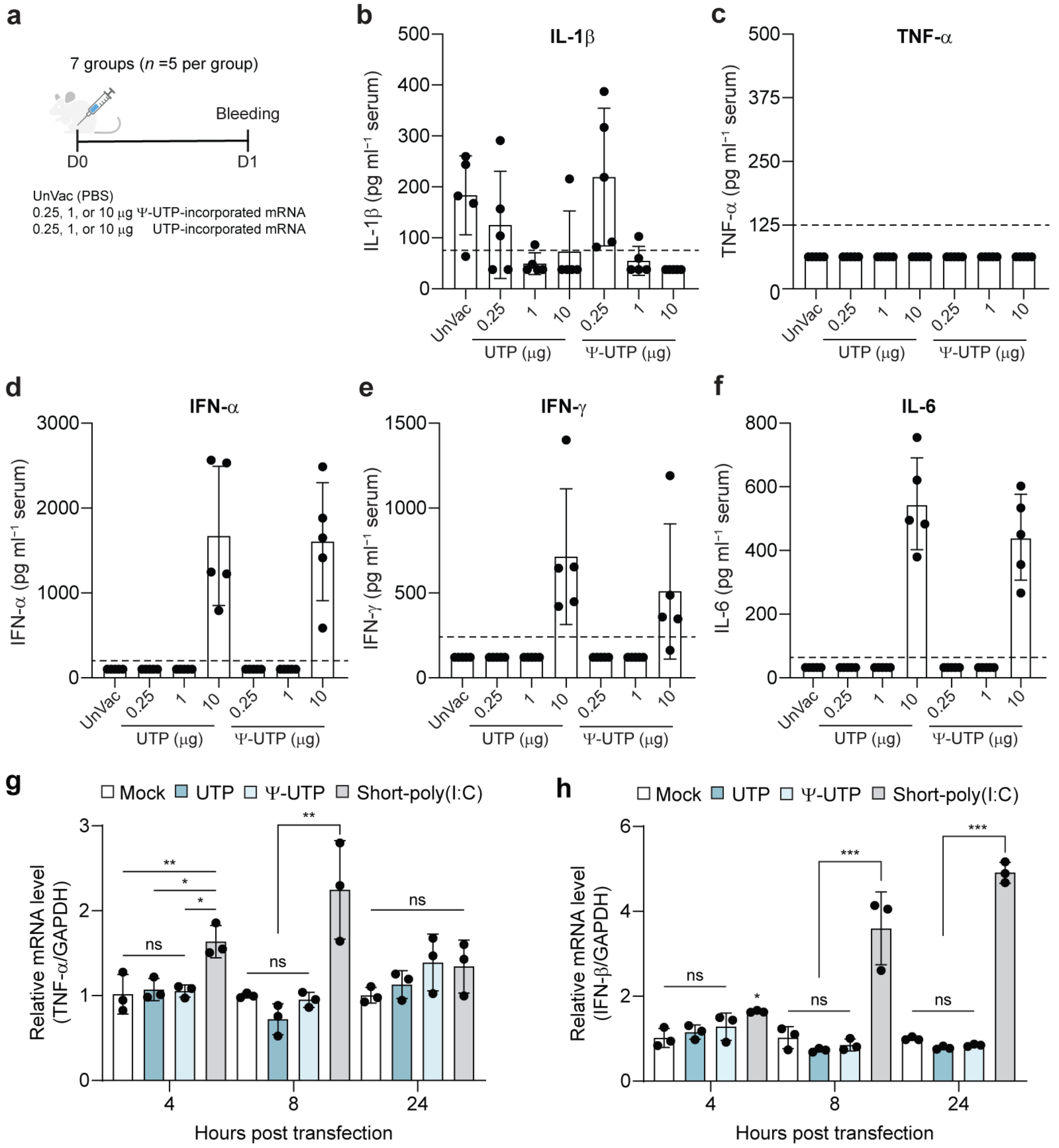


**Extended Data Fig. 6. Innate immune responses to the UTP- and ψ-UTP-incorporated BA.5 S(SA) mRNA vaccines. a**, Schematic illustrating the immunization and sample collection schedule. **b–f**, Serum cytokine levels measured by ELISAs. The dotted line represents the detection limit (LOD) of each assay kit. Values below the LOD were plotted at half of the LOD value for graphical representation. Each data point represents values from two technical duplicates. **g–h**, Inflammatory innate immune responses induced by mRNA-LNP transfection (0.5 μg) in HEK293T cells, assessed by RT-qPCR for the indicated cytokines. Each data point represents values from two technical duplicates. Statistical significance was determined using an unpaired Student *t*-test. * $P$<0.05; ** $P$ <0.01; *** $P$ < 0.001; ns, not significant.

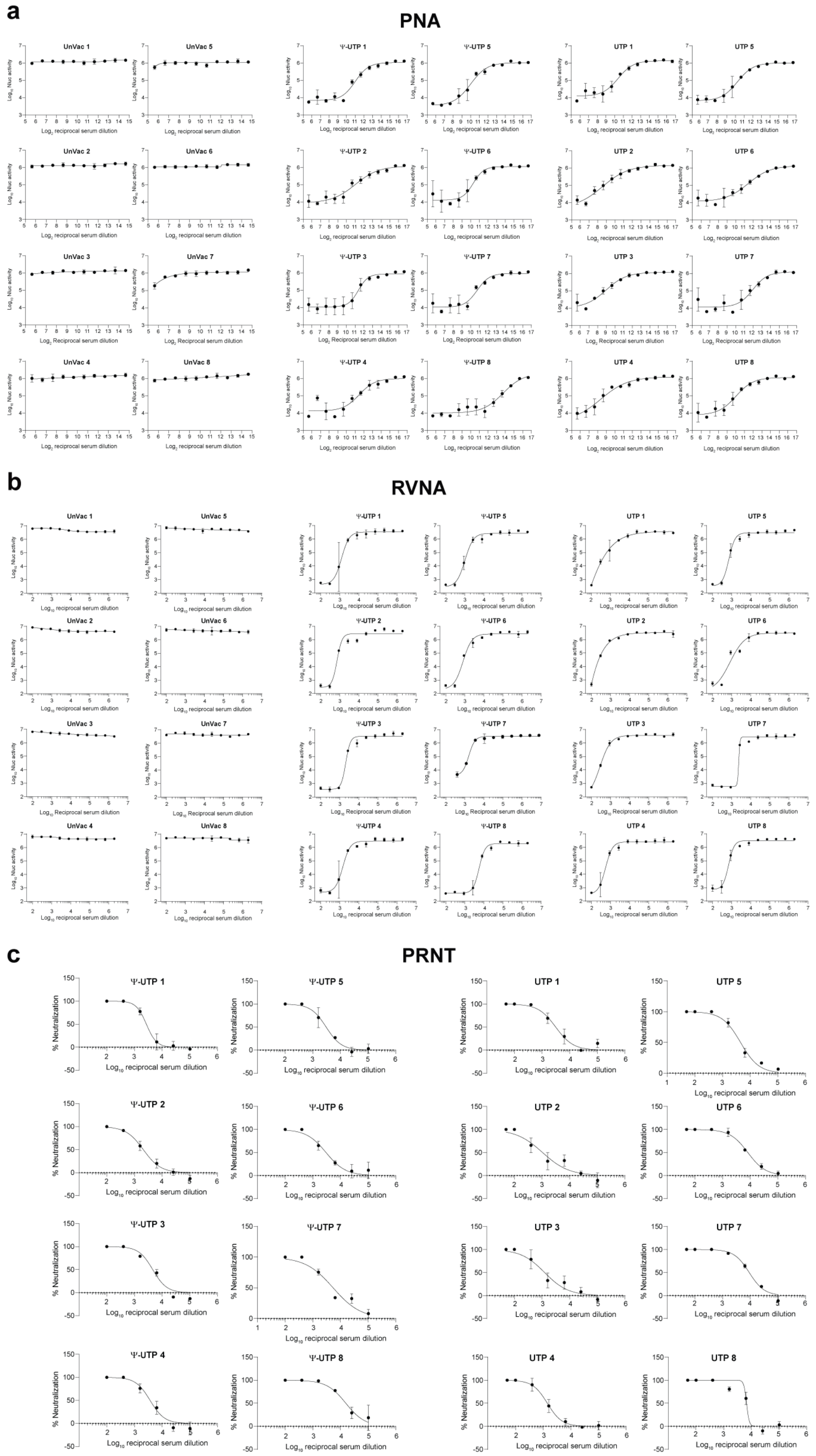


**Extended Data Fig. 7. Neutralizing antibody titers against the Omicron BA.5. a**, Neutralizing IgG levels against Omicron BA.5 spike-loaded pseudovirus were titrated using the pseudovirus neutralization assay (PNA). $Log_{10}$ ALU vs.

– continued next page

$log_2$ reciprocal serum dilution fold was plotted for each sample. Each data point represents $log_{10}$ ALU from the biological triplicates of each sample. **b**, Reporter virus neutralization assay (RVNA) was carried out to determine neutralizing IgG levels against Omicron BA.5 S- and reporter (Nluc)-expressing recombinant SARS-CoV-2, rYS006(BA.5_S)_Nluc. $Log_{10}$ ALU vs. $log_{10}$ reciprocal serum dilution fold was plotted for each sample. Each data point represents $log_{10}$ ALU from the biological duplicate of each sample. **c**, Neutralizing IgG levels against SARS-CoV-2 BA.5 variant were titrated using the plaque reduction neutralization test (PRNT). Percentage neutralization vs. $log_{10}$ reciprocal serum dilution fold was plotted for each sample. Each data point represents % neutralization from a biological duplicate of each sample.

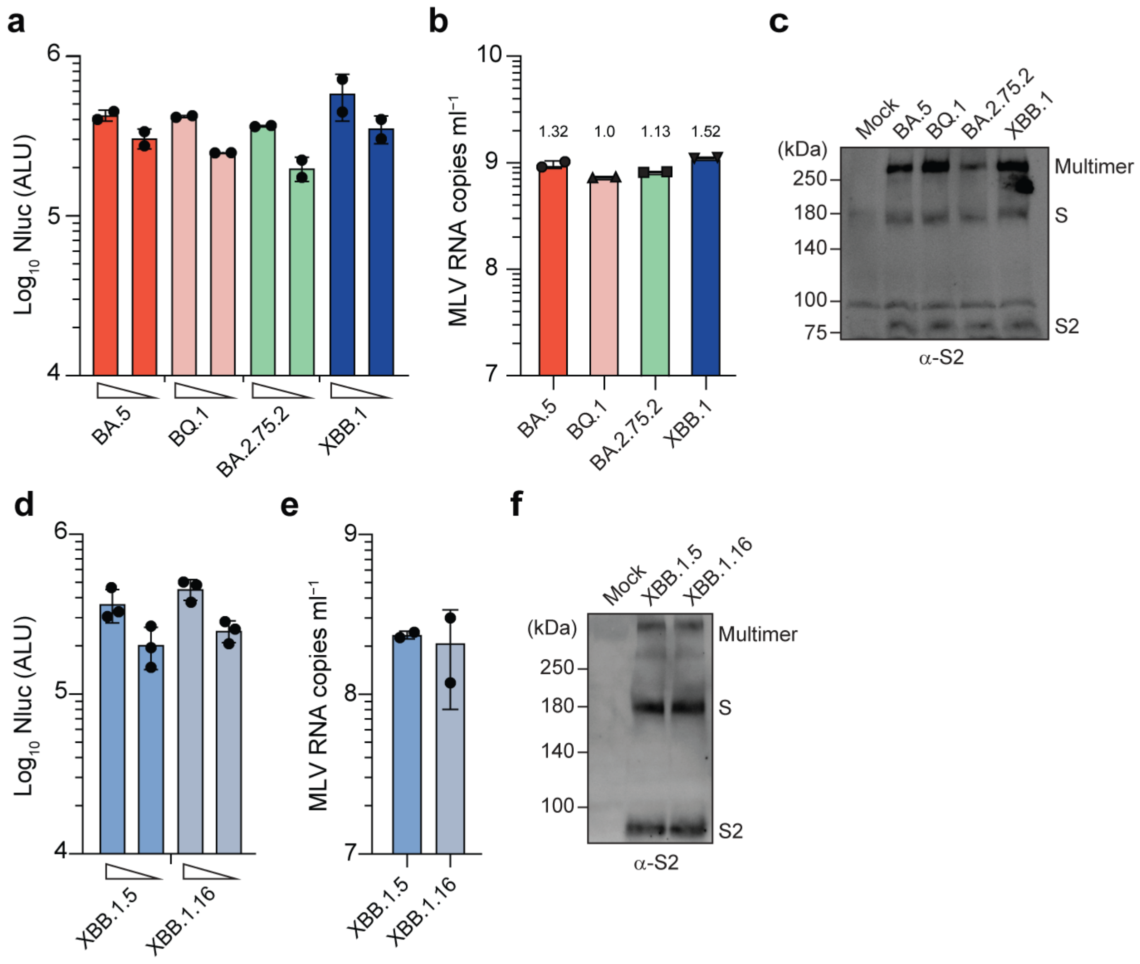


**Extended Data Fig. 8. The Omicron subvariant spike-loaded pseudoviruses used to evaluate the cross-reactivity of BA.5 S(SA) mRNA vaccines.** The pseudoviruses loaded with the S variants of Omicron subvariants were analyzed to use an equal titer of the viruses for neutralizing assay. **a**, The luminescence (Nluc) levels in Vero E6/A2-T2 cells transduced with two different doses of the pseudovirus preparations. **b**, MLV genomic RNA levels, determined by RT-qPCR using the MLV LTR in vitro transcripts as a standard, in the pseudovirus stocks (normalized to the Nluc activity). **c**, The S variant levels of the pseudovirus stocks (normalized to MLV genome copy number) used in the cross-reactivity tests were analyzed by immunoblotting. **b**–**f**, Analysis of the XBB.1.5 or XBB.1.16 S protein-loaded pseudoviruses, as described in **a**–**c**.

**Supplementary Table 1. Profile of the S variants with a single amino acid change in VeroE6 culture-adapted NCCP 43326 stocks**

| Passage no | Nucleotide position[a] | Total read depth | Read no. w/ nt change[b] | Read no. w/o nt change | Proportion of variant (%) | Amino acid substitution (nt change) |
|---|---|---|---|---|---|---|
| P1 | 1963 | 195 | 39 | 156 | 20.0 | H655Y (C1963T) |
| | 2044 | 181 | 10 | 169 | 5.52 | R682W (C2044T) |
| | 2053 | 184 | 85 | 99 | 46.2 | R685S (C2053A) |
| | 2054 | 188 | 8 | 179 | 4.26 | R685H (G2054A) |
| P2 | 2053 | 195 | 169 | 26 | 86.7 | R685S (C2053A) |

[a]Nucleotide positions are numbered according to the spike protein-coding gene sequence of the reference strain, Wuhan-Hu-1 ((hCoV-19/Wuhan/Hu-1/2019, GenBank accession no, MN908947).
[b]Read depths with a value of ≤ 2 were regarded as sequencing errors and consequently were excluded from the count.

**Supplementary Table 2. Profile of the S variants with deletions in VeroE6 culture-adapted NCCP 43326 stocks**

| Passage no | Nucleotide position[a] | Total read depth | Read no w/ nt deletion[b] | Read no w/o nt change[c] | Proportion of variant (%) | Amino acid deletions |
|---|---|---|---|---|---|---|
| P1 | 2023-2031 | 179 | 13 | 163 | 7.26 | $\Delta QTQ_{675\text{-}677}$ |
| | 2021-2035 | 181 | 3 | 169 | 1.66 | $\Delta QTQTN_{675\text{-}679}$ |
| P2 | 2023-2031 | 178 | 9 | 164 | 5.06 | $\Delta QTQ_{675\text{-}677}$ |
| | 2021-2035 | 182 | 5 | 171 | 2.75 | $\Delta QTQTN_{675\text{-}679}$ |

[a]Nucleotide positions are numbered according to the spike protein-coding gene sequence of the reference strain, Wuhan-Hu-1.

[b]Number of contigs that encompasses the nucleotide deletions leading to three or five amino acid deletions

[c]Average read depth calculated by considering the read counts for each nucleotide within the indicted nucleotide range
